# Loss of the Y chromosome drives basal identity and metastatic potential via epigenetic reprogramming in pancreatic cancer

**DOI:** 10.64898/2026.10.06.756936

**Authors:** Madeleine Dorsch, Felix E. G. Beaudry, Anna K. Baginska, Nancy Meyer, Anne Petzold, Peihua Zhao, Belen Otero-Carrasco, Kazeera Aliar, Sebastian Urban, Julia Zummack, Jennifer Jung, Johanna M. Hölken, Ningjun Duan, Laura Godfrey, Mei-Ju May Chen, Kathleen Schlüter, Katyayni Ganesan, Florian Over, Thomas J. Y. Kono, Hans-Ulrich Schildhaus, Katja Steiger, Roland F. Schwarz, Elisabeth Hessmann, Günter Schneider, Jens T. Siveke, John P. Neoptolemos, Teresa Peccerella, Maria Llamazares-Prada, Alexander Schramm, Christoph Plass, Florian Rambow, Faiyaz Notta, Barbara T. Grünwald, Barbara M. Grüner

## Abstract

Loss of the Y chromosome (LOY) has been largely unexplored in pancreatic ductal adenocarcinoma (PDAC), and no mechanistic framework previously explained its role in tumor biology. We identify LOY as a recurrent, functionally consequential alteration affecting ∼50% of male PDAC patients across two independent cohorts, with strong enrichment in advanced disease and, critically, in the basal subtype, establishing LOY as a clinically actionable biomarker. Comprehensive genomic profiling positions LOY as a distinct genetic event associated with TP53 mutations, implicating it in aggressive tumor evolution. Integrative analyses across bulk tumors, single-cell datasets and CRISPR-engineered PDAC models converge on epithelial-to-mesenchymal transition (EMT) as the dominant LOY-driven program. Y-chromosome deletion alone induces EMT transcriptional states, mesenchymal morphology and enhanced metastatic ability. Mechanistically, LOY disrupts epigenetic regulation through loss of Y-linked chromatin modifiers, reshaping DNA methylation at EMT- and basal-defining loci and reducing chemotherapy sensitivity. These findings establish LOY as a causal genetic driver of PDAC progression and a clinically accessible biomarker with direct therapeutic relevance.

## MAIN / INTRODUCTION

Loss of the Y chromosome (LOY) in blood cells represents the most frequent post-zygotic aneuploidy in males and has emerged as a major focus in biomedical research, with intensive efforts now directed toward dissecting its link to aggressive tumor phenotypes^1,2^. Once regarded as a benign, age-related anomaly, mosaic LOY (mLOY) is now recognized as a pervasive somatic event with profound biological consequences^3–5^. Its prevalence increases exponentially with age, affecting ∼20% of men by the age of 80 and exceeding 57% in those over 90 years^6,7^. Environmental exposures such as smoking and genetic predisposition further accelerate its occurrence, particularly in proliferative compartments like hematopoietic cells^4,8–10^. Accumulating evidence links mLOY to immune dysfunction and heightened susceptibility to neurodegenerative, cardiovascular, infectious, and malignant diseases^7,11–14^. Mechanistically, mLOY is thought to arise from mitotic segregation errors, generating dynamic clonal expansions that reshape genomic balance^1,15^. Beyond blood, loss of the Y chromosome is also frequently observed in cancer cells across many tumor types, with prevalence ranging from very low to over 80% depending on the cancer context^12^. In tumors, LOY is typically inferred from reduced Y gene expression or decreased Y chromosome read depth in sequencing data. Importantly, tumor-associated LOY is not correlated with patient age, suggesting distinct biological mechanisms from hematopoietic mLOY^12,16^. LOY has been implicated across multiple cancer entities including bladder, prostate, kidney, colorectal, lung, and esophageal tumors where it correlates with aggressive phenotypes, immune exhaustion, and poor clinical outcomes^12,14,17–20^. Several Y-linked genes with tumor-suppressive functions, such as *KDM5D, UTY, EIF1AY* and *DDX3Y,* are consistently downregulated in LOY cancers, and have been implicated in important processes such as cell cycle regulation, tumor progression, and immune evasion^12,21–23^.

Despite these insights, the biological and clinical significance of LOY in pancreatic ductal adenocarcinoma (PDAC), one of the most lethal malignancies worldwide, is mostly unclear. Metastatic spread is the principal cause of mortality in PDAC, yet, unlike tumor initiation, no single genetic mutation has been identified as a universal driver of metastasis^24,25^. Preliminary studies confirm the presence of LOY in PDAC, but its biological and clinical significance, particularly in shaping tumor plasticity, progression, and metastasis within a disease marked by extreme plasticity, early dissemination, and dismal survival, remains unexplored^12,14,26^.

Here, we demonstrate that LOY is not a bystander but an active driver of epithelial-to-mesenchymal transition (EMT) and the more aggressive basal subtype in PDAC. Through integrative molecular and functional analyses, we reveal that LOY reprograms transcriptional networks via epigenetic remodeling to promote mesenchymal-like states, thereby enhancing tumor plasticity, invasiveness, metastatic potential, and resistance to standard of care therapy. These findings establish LOY as a mechanistic contributor to PDAC progression, positioning it as both a potential biomarker for patient stratification and a source of novel therapeutic vulnerabilities. By uncovering increased EMT signaling and the basal subtype associated with LOY cells, our study opens new avenues for translational intervention in one of the most intractable cancers.

## RESULTS

To assess whether LOY represents a critical determinant in PDAC, we analyzed 239 male patients from the Toronto COMPASS cohort (241 patient samples, 147 male patient samples), and a cohort of the resected PDAC cases (168 patient samples, 92 male patient cases)^27^ of which whole-genome data from laser-capture-microdissected (LCM)-tumor tissue were available, excluding cases with low tumor content (≤ 30%, n = 13). Stratification of Y chromosome copy numbers across male patients revealed a clear separation between LOY (n = 106) and WTY (wild-type Y, n = 95) cases, with partial Y losses (n = 38) excluded in subsequent analyses to ensure stringency and focus on complete Y loss (Fig. 1a). Firstly, neither age nor tumor cellularity differed significantly between LOY and WTY (Suppl. Fig. 1a-b). Next, comparison of large copy number alterations outside of the sex chromosomes uncovered no consistent differences between LOY and WTY cases (Fig. 1b). LOY is considered to be increased due to chromosomal instability^5^, yet no differences in structural variants could be detected between LOY and WTY (p = 0.52) (Suppl. Fig. 1c). However, comparing the frequency of recurrent PDAC driver mutations between LOY versus WTY revealed a significantly higher frequency of *TP53* mutations in LOY patients (LOD = -1.57 [CI = -2.57:-0.685]; p = 0.0001). Furthermore, we observed a significant increase of mutations in *KDM6A* in LOY (LOD = -2.43 [-4.64:-0.98]; p = 7e-05) and *TGFBR2* in WTY patients (LOD = 1.21 [0.16-2.40]; p = 0.01), respectively (Fig. 1c). However, when we compared for *KRAS* imbalance, defined as having more copies of the mutant variant than of the wildtype^28^, between LOY and WTY patients, we did not detect a significantly different frequency of *KRAS* imbalance in LOY patients (n = 181, p = 0.18, Fig. 1d).

**Figure 1.**
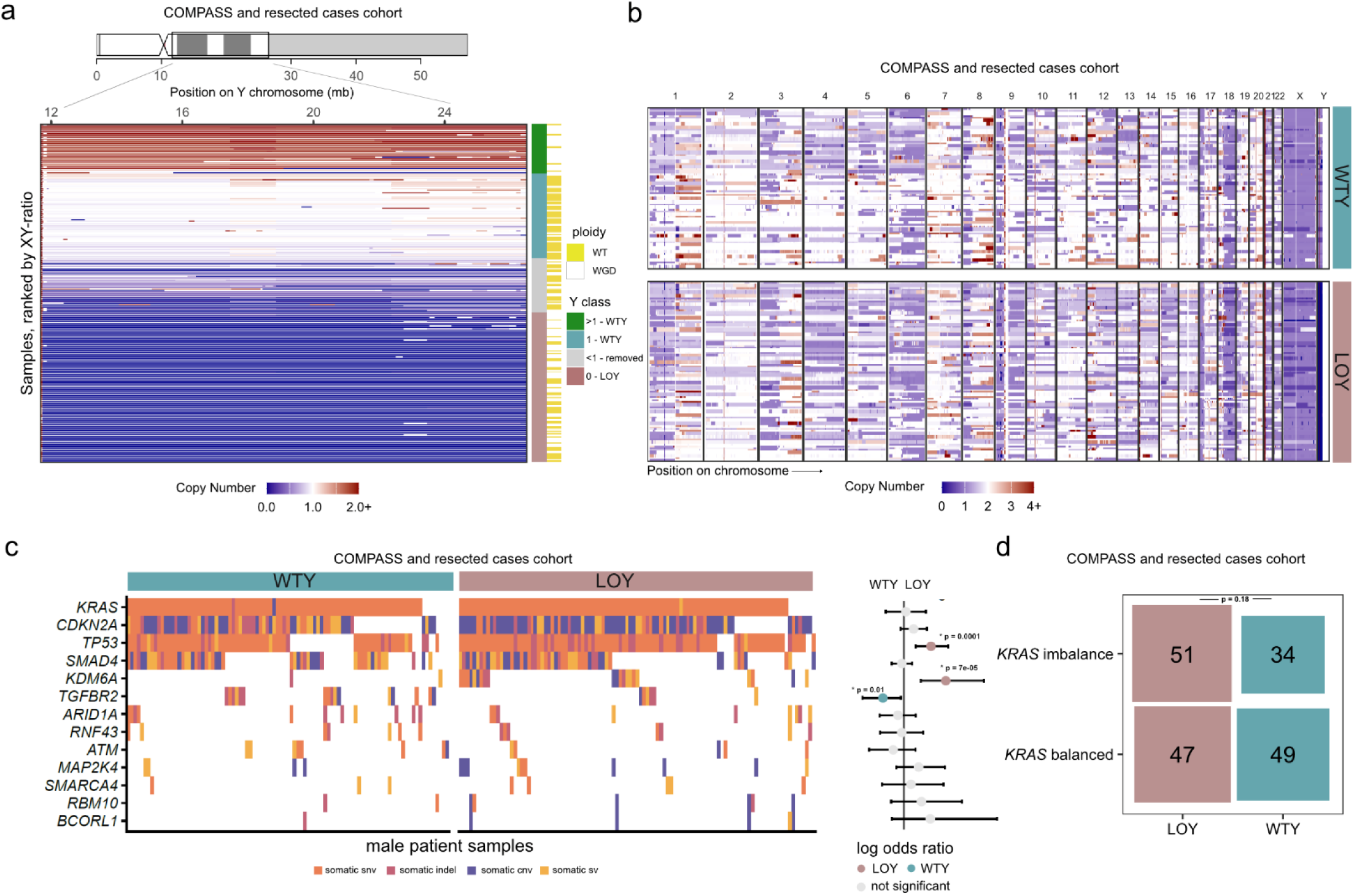
Genomic consequences of LOY in male PDAC patients. **a)** Copy number profiles of whole genome sequencing across the Y chromosome of all male patients (COMPASS and resected PDAC cases cohort). Ploidy was classified as wildtype (WT) or whole-genome duplicated (WGD). Y-chromosome status was defined as wildtype Y (WTY) or loss of Y (LOY). **b)** Copy number profiles of whole-genome sequencing across the entire genome of all male patients (COMPASS and resected PDAC cases cohort). **c)** Distribution of the most common PDAC somatic alterations, stratified by LOY and WTY status and associated log odds ratio. **d)** Association between LOY and *KRAS* allelic imbalance. Box plots compare the counts of *KRAS* imbalance versus balanced cases between LOY and WTY samples.

To systematically assess whether LOY exerts direct functional effects in pancreatic cancer, we deleted the entire Y chromosome in male PDAC cell lines. We employed a previously established CRISPR/Cas9 strategy that generates multiple double-strand breaks across Y-specific repetitive regions^29^. Guide RNAs targeting highly repeated loci on the short arm, long arm, and centromere of the Y chromosome were used to induce extensive fragmentation and promote chromosome loss during subsequent cell divisions. Interestingly, of all male PDAC cell line models available on Cellosaurus^30^, over 75% already lacked the amelogenin short tandem repeat (STR) Y-profile marker (Fig. 2a), therefore displaying at least partial loss of the Y chromosome. Furthermore, both methods FISH for the long as well as the centromer of the Y chromosome and Y chromosome specific PCR^31^ confirmed that even some STR-positive cell lines carried only partial Y chromosome. For example, CF-Pac1 cells retained only the short arm Yp (Suppl. Fig. 2a-c). Among the initially tested models, YAPC and HuP-T3 cells harbored an intact Y chromosome, validated by FISH or Y chromosome-specific PCR (Fig. 2b, Suppl. Fig. 2a, b, d-f). Therefore, from both YAPC and HuP-T3 cells, several independent clones with complete Y loss were generated using the CRISPR/Cas9 strategy, and each four YKO clones were selected for further characterization (Fig. 2b, Suppl. Fig. 2d-f). First, we performed whole-genome sequencing of two selected clones YAPC-YKO-1, YAPC-YKO-3, and the parental YAPC line and observed that, aside from the complete Y loss (Fig. 2c, Suppl. Fig. 2g), no additional copy-number alterations were detected. This confirms the specificity of the CRISPR approach, excludes clonal selection artifacts, and establishes a clean genetic background for downstream functional analyses. Moreover, we established that FISH and Y chromosome–specific PCR serve as robust, low-complexity alternatives to whole-genome sequencing for confirming Y loss status.

**Figure 2.**
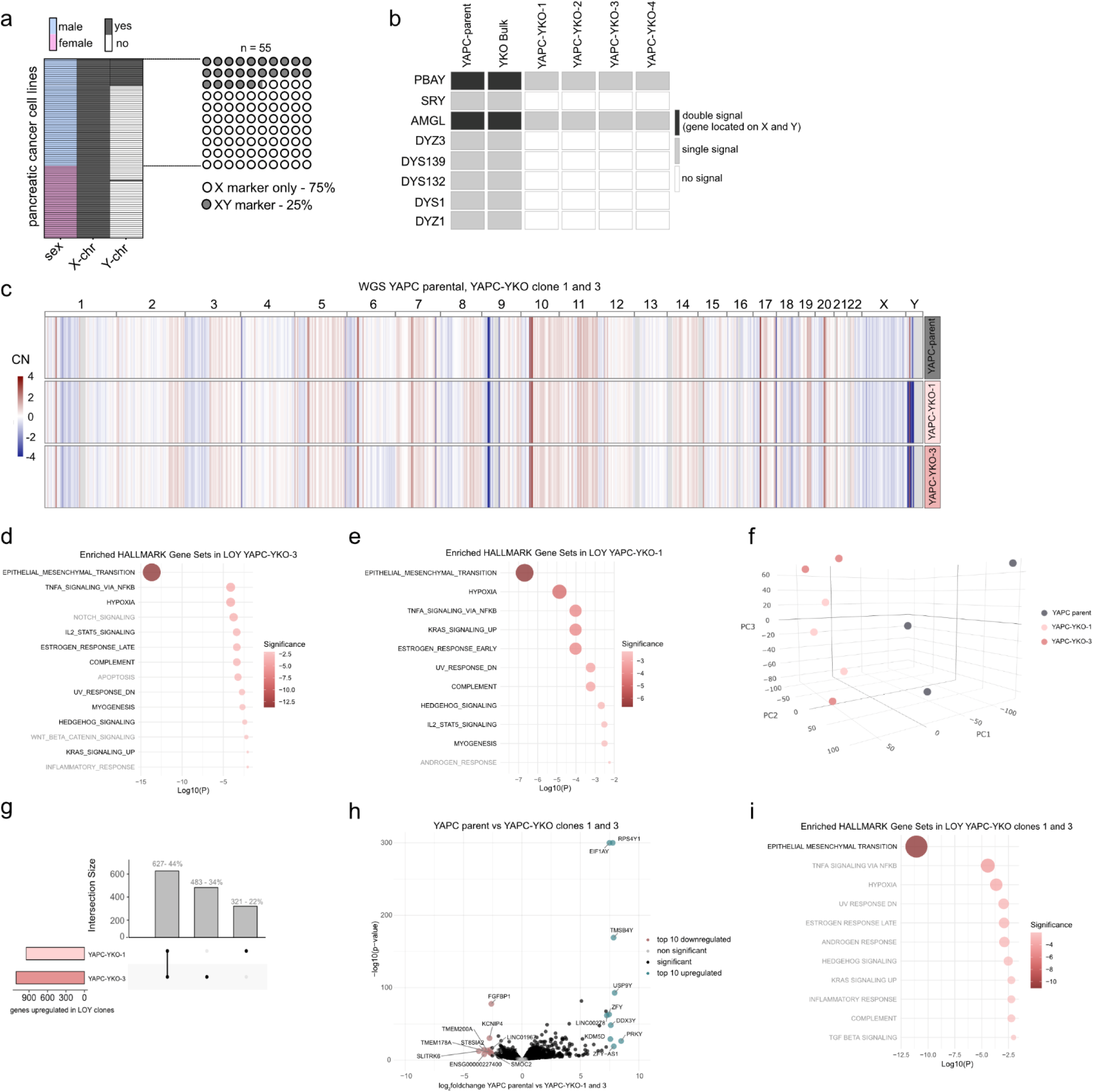
Genomic and transcriptomic characterization of functional CRISPR-CAS-9-Knockout of the Y chromosome in PDAC cell lines. **a)** STR marker analysis of available human pancreatic cancer cell line models using the Cellosaurus database (X/Y status assessed via Amelogenin profiling). **b)** Semi-quantitative analysis of Y chromosome PCR. Table summarizing signal detection for nine Y-linked genes across YAPC-YKO clones. AMGL and PBAY show X- and Y-derived PCR products, all other assayed genes are exclusively Y-linked and yield a single PCR product. **c)** Genome-wide CNV profiles from whole-genome sequencing of YAPC parent cells and YAPC-YKO-1 and YAPC-YKO-3 clones. Heatmap of GC-corrected log_2_ copy number ratios calculated from 20 kb genomic bins for three WGS samples. Genomic bins are ordered by chromosomal position and annotated by chromosome (top). Copy number losses and gains are shown in blue and red, respectively. **d, e)** Pathway enrichment analysis of RNA-seq data comparing YAPC-YKO 1 (d) and YAPC-YKO 3 (e) clones to YAPC parental cells. Dot plots display significantly enriched HALLMARK pathways among genes upregulated in YKO clones (log_2_ fold change < –0.75, adjusted p < 0.05). **f)** Principal component analysis (PCA). 3D PCA of transcriptomes from RNA-Seq analysis of YAPC parent and YAPC-YKO-1 and YAPC-YKO-3 clones. **g)** Intersection of significant, differentially expressed genes between YKO clones 1 and 3 (log_2_ fold change < –0.25, adjusted p < 0.05). **h)** Volcano plot of differential gene expression analysis comparing YAPC-parent to YAPC-YKO-1 and YAPC-YKO-3 clones. **i)** Pathway enrichment analysis of RNA-seq data comparing YAPC-YKO-1 and YAPC-YKO-3 clones combined to YAPC-parent cells. Dot plot shows significantly enriched HALLMARK pathways from genes upregulated in YKO clones.

To analyze the impact of the Y chromosome loss on transcriptional regulation, we performed RNA-seq analysis on the YAPC parental cells and YAPC-YKO-1 and YAPC-YKO-3 clones to complement the WGS dataset and ensure a high-quality, genetically matched dataset rather than introducing variability through additional clones. Notably, with focus on Hallmark pathways only, EMT emerged as the top enriched pathway in both LOY clones when compared to parental YAPC cells (Fig. 2d, e). Overall, pathway signatures of both up- and downregulated genes were highly similar across both YAPC-YKO-1 and YAPC-YKO-3, and principal component analysis (PCA) confirmed tight clustering with substantial overlap in up- or downregulated genes and pathways (Fig. 2f, g; Suppl. Fig. 2h,i). Consequently, to robustly visualize LOY-associated transcriptional changes and minimize clone-specific effects, YAPC-YKO-1 and YAPC-YKO-3 were combined into a unified LOY group for downstream analyses. As expected, Y-linked genes were strongly upregulated in parental cells (Fig. 2h), whereas EMT emerged as the most significantly enriched pathway in YAPC-YKO-1 and YAPC-YKO-3 (Fig. 2i). Downregulated pathways also showed a high degree of overlap (Suppl. Fig. 2h-j).

All YKO clones displayed marked morphological changes consistent with a more mesenchymal-like phenotype, which remained stable upon passaging in both 2D and 3D culture, whereas parent YAPC and HuP-T3 cells presented a more epithelial-like morphology (Fig. 3a, Suppl. Fig. 3a,b). Reflecting this, mesenchymal markers such as *VIMENTIN* and *SNAIL2* were upregulated in both cell line models across most clones (Fig. 3b-c, Suppl. Fig. 3e-f). Also, YAPC-YKO clones displayed reduced colony-forming capacity, a characteristic of mesenchymal-like cells that fail to establish robust epithelial networks (Suppl. Fig. 3c,d). Consistently, anoikis survival assays, which provide a direct functional measure of a mesenchymal-like cell state *in vitro*^32^, demonstrated that YKO clones from both cell lines (YAPC and HuP-T3) were significantly more protected against anoikis-induced cell death (Fig. 3d; Suppl. Fig. 3g,h,j).

**Figure 3.**
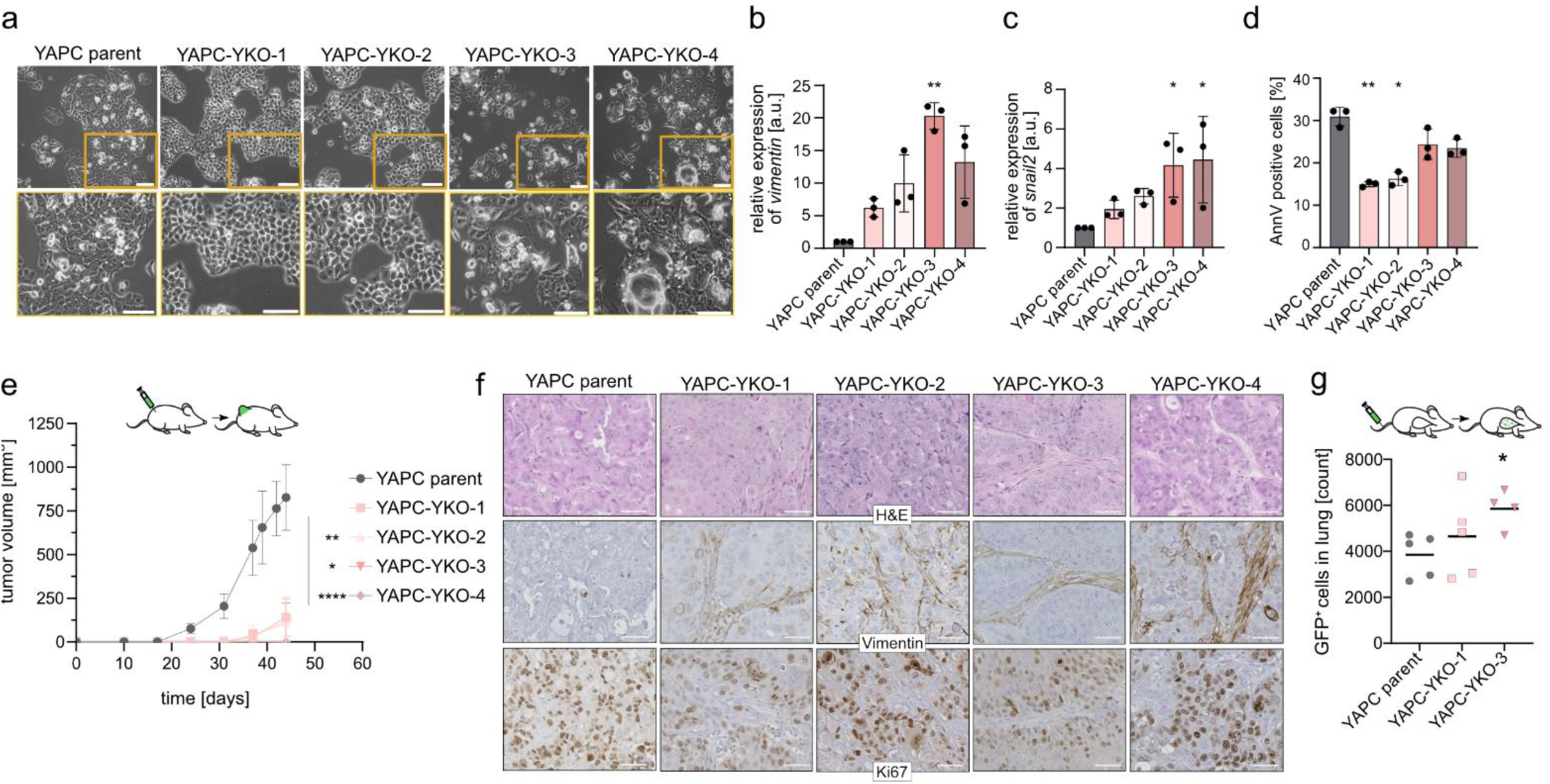
Functional *in vitro* and *in vivo* analysis of YKO cell line clones. **a)** Representative 2D morphology images of YAPC-parent cells and all four YAPC-YKO clones. Magnified regions highlighted by yellow boxes. Scale bars represent 100 µM. **b, c)** Quantitative RT-PCR analysis of mesenchymal markers *Vimentin* (b) and *SNAIL2* (c), normalized to *GAPDH* in YAPC-parent versus YAPC-YKO clones. Three biological replicates are displayed. Statistical significance was assessed using the Kruskal–Wallis test. **d)** Quantification of cells undergoing anoikis in YAPC-parent and YAPC-YKO clones. One representative out of three independent experiments is depicted. Statistical significance was assessed using the Kruskal–Wallis test. **e)** Subcutaneous tumor growth of YAPC-parent cells and YAPC-YKO clones injected into NSG mice to ascertain initial tumor formation capacity *in vivo*. Statistical significance was assessed using the Kruskal–Wallis test. **f)** Histological and immunohistochemical analysis of subcutaneous tumors as shown in E). Representative H&E, Vimentin, and Ki-67 staining of tumor sections from YAPC-parent and YAPC-YKO clones. Scale bars represent 50 µM. **g)** Intravenous injection of YAPC-parent cells and YAPC-YKO-1 and YAPC-YKO-3 to ascertain survival in the bloodstream and initial seeding to lung quantified using flow cytometry (FACS) of GFP-positive cells in the lungs. Statistical significance was assessed using the Kruskal– Wallis test.

Strikingly, we could confirm a profound mesenchymal-like cell state in YKO clones also *in vivo*. Following subcutaneous injection into NSG mice, all four YAPC-YKO clones exhibited a drastically reduced ability to initiate tumor formation, a hallmark of mesenchymal cells that cannot undergo mesenchymal-to-epithelial transition (MET) to establish epithelial tumor masses (Fig. 3e)^33^. Histological analyses confirmed pronounced morphological alterations from either a solid or ductal morphology towards a squamous phenotype and a strong increase in Vimentin expression within tumor stroma, while Ki67 levels remained comparable to controls once tumors formed (Fig. 3f). These findings demonstrate that the delayed tumor growth was not due to impaired proliferation, but rather to a loss of epithelial tumor-forming capacity. Conversely, intravenous injection into mice produced the opposite effect: YAPC-YKO-1 and YAPC-YKO-3 clones exhibited markedly increased initial seeding capacity to the lung, fully consistent with a mesenchymal-like phenotype where tumor cells in the initial steps of the metastatic cascade have to display mesenchymal behavior to survive, travel through the bloodstream, and adhere at a secondary site (Fig. 3g, Suppl. Fig. 3i,j)^33,34^. Overall, these results confirm at a functional level that loss of the Y chromosome alone is sufficient to induce EMT and confer a more aggressive mesenchymal phenotype in PDAC cells.

Importantly, gene set enrichment analysis using transcriptomic data of the COMPASS and resected PDAC patient cohort (n = 191)^27^ confirmed EMT signaling as one of the most significantly upregulated signaling pathways also in LOY tumors (Fig. 4a). Of note, Y chromosome encoded genes as genes located on the Y chromosome were most prominently upregulated in WTY patients (Suppl. Fig. 4a). We could also confirm a strong link of LOY in immune cells with immune evasion and cancer progression, which had been previously reported in bladder cancer^14,21^ and other cancer entities^14^. Comparison of the differentially regulated pathways from the COMPASS cohort and our *in vitro* data (Suppl. Fig. 2h-j) suggested that inflammatory responses were downregulated in the YKO clones. Together, this suggested that these transcriptomic changes in immunoregulatory pathways were not solely driven by tumor cells.

**Figure 4.**
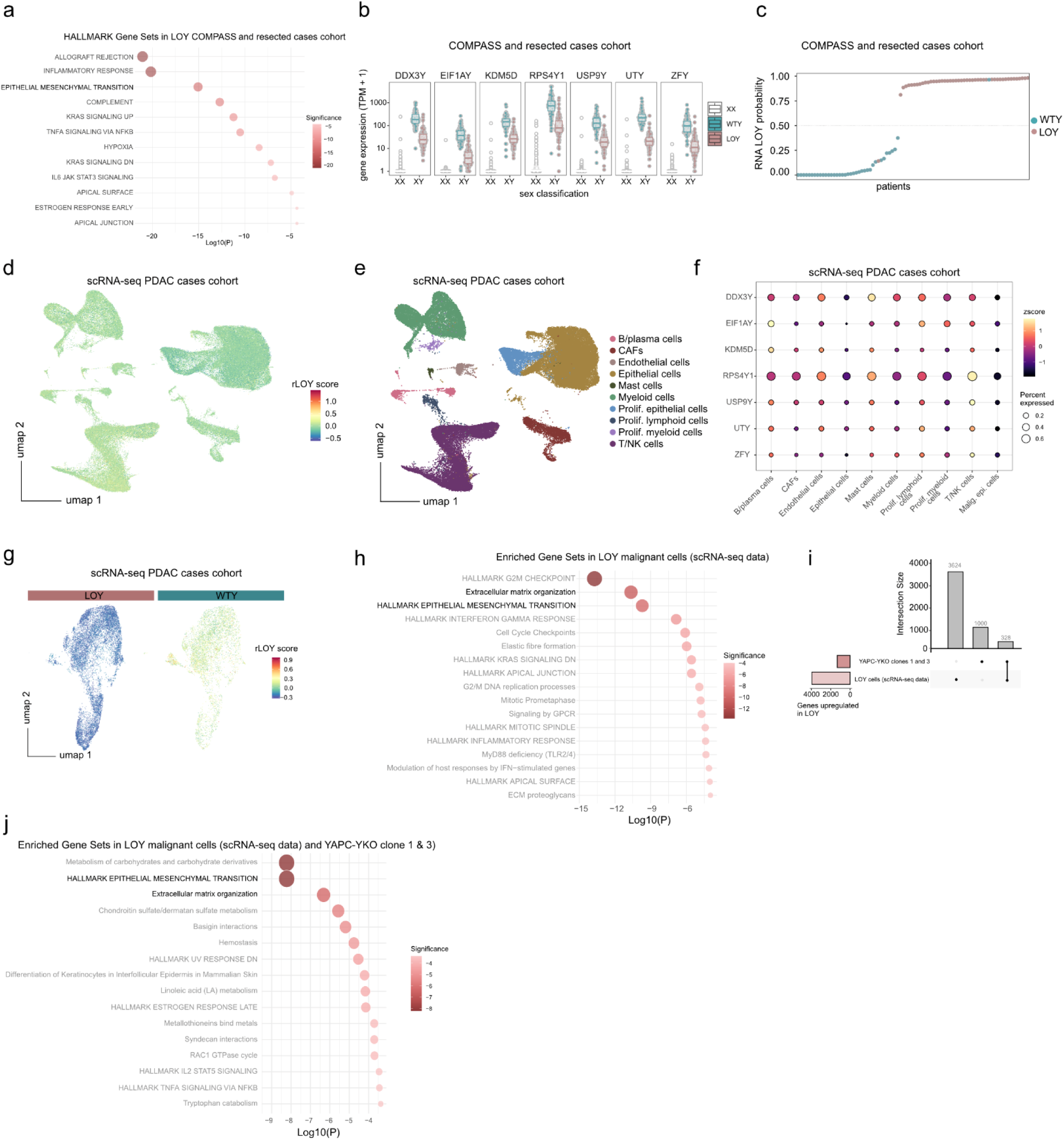
Transcriptomic profiling of LOY in two independent male PDAC patient cohorts. **a)** Pathway enrichment analysis of upregulated genes in LOY patient samples in the COMPASS and resected patient cohort. Dot plot of enriched HALLMARK gene sets in LOY samples (log_2_ fold change >0.4, adjusted p < 0.05). **b)** Relative gene expression analysis of the seven top regulated Y chromosome-encoded genes classified from WGS sex classification (XX, WTY and LOY) from the COMPASS and resected PDAC cases cohort. **c)** RNA probability score of Y chromosome gene signature expression (rLOY) across COMPASS and resected PDAC patients and correlation to WGS sex classification call. **d)** UMAP of rLOY score expression (DDX3Y, EIF1AY, KDM5D, RPS4Y1, USP9Y, UTY, ZFY) across all male patients from the Werba et al. single cell PDAC cohort. **e)** UMAP of cell type annotations across all male patients from the Werba et al. single cell PDAC cohort. **f)** Relative gene expression of the depicted 7 top Y chromosome encoded genes across all cell types from all male patients from Werba et al. single cell PDAC cohort. Dot plot visualizes expression of Y-linked genes across annotated cell types. Dot size reflects percent of expressing cells; color intensity indicates scaled expression (Z-score). **g)** UMAP of Y gene signature expression in WTY versus LOY epithelial malignant cells stratified by CopyKAT. **h)** Pathway enrichment analysis of upregulated genes in the Werba et al. single cell dataset. Dot plot of enriched HALLMARK and Reactome gene sets in rLOY malignant epithelial cells (log_2_ fold change > 0.75, adjusted p < 0.05). **i)** Intersection of significant differentially expressed genes between YAPC-YKO-1 and YAPC-YKO-3 clones combined and the upregulated genes in LOY epithelial malignant cells of the single-cell RNA patient dataset of Werba et al (log_2_ fold change < –0.25 (YAPC-YKO-1 and YAPC-YKO-3), > 0.25 (single-cell RNA patient dataset of Werba et al.), adjusted p < 0.05). **j)** Pathway enrichment analysis of overlapping genes (I) between RNA-seq data of upregulated genes in the YAPC-YKO-1 and YAPC-YKO-3 clones combined and the upregulated genes of the LOY epithelial malignant cells from the sc-RNA-seq patient dataset of Werba et al. Dot plot shows significantly enriched HALLMARK and Reactome pathways from genes upregulated in all LOY cells.

To clarify conclusively whether and how LOY in pancreatic cancer cells might regulate immunoregulatory and tumor cell signaling pathways, we analyzed a previously published single-cell dataset of PDAC patients^35^. First, we plotted the relative expression levels of seven Y chromosome encoded genes (*DDX3Y, EIF1AY, KDM5D, RPS4Y1, USP9Y, UTY* - also known as *KDM6C -,* and *ZFY*) that were established^12^ to determine LOY and that were most significantly differentially expressed in LOY versus WTY patients in the transcriptomic data of the COMPASS cohort and the resected PDAC cases (Suppl. Fig 4A). This analysis revealed a clear distinction between XX (female cases, no expression), XY (male WTY, strong expression), and LOY (male, reduced expression) genotypes (Fig. 4b). From these seven genes, we calculated an RNA-based LOY probability score (hereafter rLOY score) and found that the rLOY score was consistent with the WGS analysis in 93% of our testing set (Fig. 4c). We next applied the rLOY score to an independent scRNA-seq PDAC patient dataset^35^. Overall, rLOY varied substantially across cell types, being most prominently downregulated in epithelial cells but also showing some variation in Y-gene expression levels in cells of the tumor microenvironment (TME) (Fig. 4d-f). As our focus was on tumor cell intrinsic effects of LOY, we then subset epithelial malignant cells - i.e. cancer cells - identified as aneuploid using CopyKAT^36^ (Fig. 4g, Suppl. Fig. 4b). Overall UMAP distributions did not distinguish between

LOY and WTY cancer cells (Fig. 4g). However, differential gene expression analysis revealed that malignant epithelial rLOY cells upregulated multiple genes relative to WTY cells, leading to pathway enrichment in processes such as G2/M checkpoint transition, EMT, and interferon gamma response (Fig. 4h, Suppl. Fig. 4c). All differences observed between rLOY and WTY epithelial malignant cells were independent of gene number, UMI counts, or mitochondrial gene content (Suppl. Fig. 4d). Finally, we compared the transcriptome profile of YAPC-YKO-1 and YAPC-YKO-3 clones with the malignant LOY cells from the Werba et *al*. single cell dataset to focus on common regulated genes^35^. The overlap revealed 328 common differentially upregulated genes (Fig. 4i). Importantly, using only the genes overlapping in both LOY tumor cell datasets for a combined gene set enrichment analysis, EMT was the most significantly upregulated hallmark gene set (Fig. 4j). Overall, the enriched gene sets in LOY cancer cells derived both from the experimental YAPC-YKO clones and the scRNA-seq patient dataset^35^, as well as the results from the COMPASS and resected PDAC patient cohorts matched particularly in EMT upregulation, highlighting a functional upregulation of EMT upon LOY.

Previously, a strong link between (partial) EMT and the basal, more aggressive subtype in PDAC has been established^37,38^. To examine whether LOY tumors are clinically also more basal, we applied the Moffitt-modified gene signature^39^ in the COMPASS and resected PDAC cases^27^ transcriptome data with complete clinical profiles (n = 191) and demonstrated significant enrichment of samples of the basal subtype in the LOY group compared to classical samples (p = 0.0001613). The frequency of LOY further increased with advanced tumor stages (Fig. 5a, b, Suppl. Fig. 5a). Unsupervised clustering of basal and classical genes confirmed the association of LOY with the basal subtype (LOY odds ratio 2.62 [1.40-4.92], p = 0.001) (Fig. 5c). Furthermore, differential gene expression analysis (log_2_ fold change) highlighted that 23 of 24 expressed basal signature genes were upregulated in LOY samples, whereas 24 of 25 expressed classical signature genes were upregulated in WTY tumors, underscoring the pronounced transcriptional coupling between LOY and the basal, aggressive PDAC subtype (Fig. 5d). Finally, to ensure LOY cells both belong to the basal subtype and are also enriched in EMT signature gene expression, we examined EMT gene scores from the GSEA hallmark set alongside basal and classical subtype scores defined in Figure 5D in the scRNA-seq dataset^35^. We found that LOY cancer cells were enriched for both EMT and basal programs, and that cancer cells with a stronger EMT signature were also enriched in basal gene expression, whereas WTY cancer cells displayed higher classical scores (Fig. 5e-g, Suppl. Fig. 5b). This could even be observed, to a slightly lesser degree, for the entire subset of epithelial cells without any further filtering for aneuploid/malignant cells (data not shown). We further stratified the dataset by patient, comparing patients with complete LOY versus patients with WTY or incomplete LOY (“other”) in epithelial malignant cells (Suppl. Fig. 5c). Despite patient-level heterogeneity in LOY (Suppl. Fig. 5d), LOY patients showed enrichment of G2/M checkpoint, EMT, and extracellular matrix (ECM) pathways (Suppl. Fig. 5e), with higher expression of EMT and basal genes, whereas classical genes were more highly expressed in WTY patients (Suppl. Fig. 5f-h). These results confirm a clear correlation between LOY and both EMT and the basal PDAC subtype.

**Figure 5.**
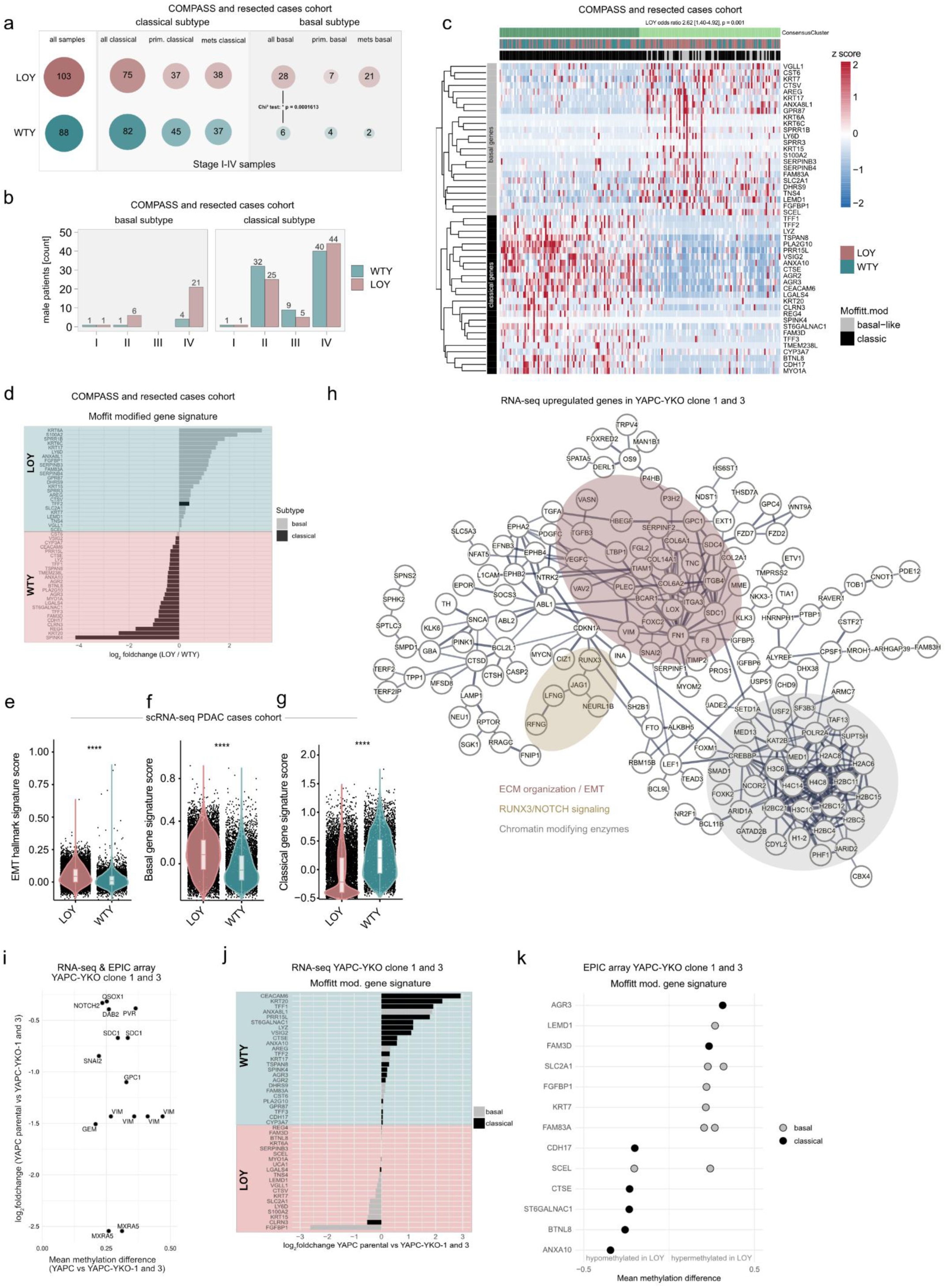
Functional connection of LOY with the basal subtype via epigenetic regulation. **a)** Distribution of LOY and WTY cases across molecular subtypes and tumor stages in the COMPASS and resected PDAC cases patient cohort. Bubble plot shows the number of LOY and WTY samples across Moffitt.mod classical and basal subtypes of primary and metastasis derived samples. Statistical significance was determined using a chi^2^-test, p = 0.0001613. **b)** LOY frequency across tumor stages within molecular subtypes. Bar charts display LOY and WTY counts across stages I–IV for basal and classical subtypes in the COMPASS and resected PDAC cases patient cohort. **c)** Gene expression consensus clustering heatmap stratified by LOY, WTY and molecular subtype according to Moffit.mod. gene signature in the COMPASS and resected PDAC cases patient cohort. Z-scored expression profiles of displayed genes are shown across LOY and WTY samples for all basal and classical genes. Statistical significance was determined using a fishers-exact test. **d)** Differential expression (log_2_ foldchange) of Moffitt-mod. signature genes in the COMPASS and resected PDAC cases patient cohort. Bar plot shows log_2_ fold changes (LOY vs. WTY) for subtype-specific genes. Genes are grouped by basal and classical annotation. **e-g)** Violin plots compare EMT hallmark (e), Moffitt.mod basal (f), and Moffitt.mod classical (g) gene signature scores between LOY and WTY epithelial malignant cells in the Werba et al. single cell dataset. Statistical significance was determined using Wilcoxon Test, **** p < 0.0001. **h)** STRING gene network analysis. Interaction map of differentially expressed genes in YAPC-YKO-1 and YAPC-YKO-3 clones compared to YAPC-parent cells with a confidence interval of 0.7 (log_2_ fold change < –0.75, adjusted p < 0.05). **i)** EPIC Array analysis of YAPC-YKO-1 and YAPC-YKO-3 clones combined versus YAPC-parent cells. Scatter plot of EMT-associated hypomethylated genes showing correlation between mean methylation difference and log_2_ fold change in expression of EMT genes. Each dot per gene visualizes a different methylation site. **j)** LOY-associated modulation of basal and classical subtype genes defining the Moffitt.mod gene signature. Bar graphs showing log_2_ foldchange expression of basal and classical subtype genes in YAPC-YKO-1 and YAPC-YKO-3 clones combined versus YAPC-parent cells from RNA-seq. **k)** EPIC array analysis of YAPC-YKO-1 and YAPC-YKO-3 clones combined versus YAPC-parent cells. Scatter plot of LOY-associated hypo- and hypermethylated genes showing mean methylation difference of Moffitt.mod basal and classical genes. Each dot per gene visualizes a different methylation site.

Finally, we sought to explore a mechanistic connection between LOY, EMT, and the basal subtype. To this end, we performed STRING network analysis of upregulated genes in our YAPC-YKO-1 and YAPC-YKO-3 clones with a focus on both Hallmark and Reactome pathways. In LOY clones, chromatin-modifying enzymes were upregulated and appeared to be linked to EMT/ECM gene programs (Fig. 5h; Suppl. Fig. 5i). Among the top enriched pathways, we identified epigenetic regulation by the WDR5 complex. WDR5 (WD40-repeat– containing protein 5) is a multifunctional chromatin regulator that plays a central role in histone modification and promotes H3K4 methylation^40^. The upregulation of this pathway is mechanistically coherent: one of the major epigenetic regulators encoded on the Y chromosome is KDM5D, a member of the KDM5 family of H3K4me2/3 demethylases that normally repress target gene expression^23,41–43^. Thus, loss of KDM5D in YKO clones removes a key H3K4 demethylase, and the compensatory upregulation of the WDR5-associated H3K4 methylation machinery represents a coherent counterpart to this deficiency. In addition to KDM5D, the Y chromosome also encodes UTY (KDM6C), a lysine demethylase that regulates H3K27 demethylation^44^. Together, these enzymes shape chromatin accessibility at lineage- and state-defining gene programs. Their combined loss in LOY clones would therefore be expected to remodel the epigenetic landscape, potentially facilitating the EMT-associated transcriptional rewiring. Leveraging that changes in histone methylation frequently occur with shifts in DNA methylation states, we used Illumina EPIC arrays to determine whether LOY-associated loss of these chromatin regulators is accompanied by genome-wide alterations in DNA methylation. Unsupervised clustering revealed a clear separation of parent YAPC WTY from YAPC-YKO-1 and YAPC-YKO-3 clones when visualizing the top 500 most hypermethylated and the top 500 most hypomethylated CpGs (ranked by mean methylation difference) (Suppl. Fig. 5j). These differential CpGs are localized predominantly to enhancer and promoter regions (Suppl. Fig. 5k). In-depth analysis of the methylation status of EMT genes that were differentially regulated between LOY and WTY cells revealed pronounced hypomethylation at multiple key regulators of EMT including *VIMENTIN*, *MRTX5*, and *SNAIL2* (Fig. 5i, Suppl. Fig. 5l), which corresponded to the observed higher RNA expression levels of these genes in the YAPC-YKO-1 and YAPC-YKO-3 clones. DNA hypomethylation is generally associated with increased gene expression due to enhanced accessibility of regulatory and transcriptional elements^45^. Consistent with this, among the 30 genes most hypomethylated in LOY, eleven (*ELAPOR1, PLCB4, PDE4B, BNC2, HEBP1, FRMD4A, TCF4, PLAGL1,*

*PPP2R2B, SCML2, PCBP2)* are already known to be involved in PDAC progression, and sixteen (*ELAPOR1, PLCB4, PDE4B, BNC2, GMDS, ZFHX4, HEBP1, FRMD4A, NRAS, TCF4, PLAGL1, PPP2R2B, TTYH3, SCML2, SETBP1, PCBP2)* are strongly implicated in metastasis and EMT (Supplementary Table S1). Overall, these findings confirm a direct mechanistic link of LOY with EMT and PDAC progression via epigenetic regulation. Importantly, in the LOY clones 11 out of 23 (including 5 with no change in log_2_ fold change) genes that define the basal PDAC subtype were upregulated and conversely, 18 out of 22 genes that define the classical PDAC subtype were downregulated (Fig. 5j). Consistent with these transcriptional shifts, 5 out of 6 basal subtype genes were hypomethylated, whereas 5 out of 7 classical subtype genes were hypermethylated in LOY clones (Fig. 5k), emphasizing the coherent connection of LOY, EMT and the basal subtype via epigenetic regulation in PDAC.

Both EMT and the basal subtype have repeatedly been associated with increased resistance to standard-of-care chemotherapy^38,39^. Based on our findings that LOY functionally connects EMT and the basal subtype, we hypothesized that LOY might serve as an easily accessible classifier to predict treatment response. To maximize translational relevance and ensure that our findings reflect patient-proximal biology, we focused our analyses on two complementary cohorts of patient-derived PDAC models. First, we analyzed a panel of 12 male primary PDAC cell lines derived either from patient-derived xenografts (PDX), which were further expanded as cell-derived xenografts (CDX), or from patient-derived organoids (PDO), which were subsequently differentiated into organoid-derived cells (ODC). Using these models, we determined LOY status from RNA-seq data using our rLOY signature (Suppl. Fig. 6a) and evaluated their sensitivity to gemcitabine, irinotecan, and Nab-Paclitaxel. Across all three agents, LOY cell lines showed a trend towards reduced response compared with WTY lines (Suppl. Fig. 6b,c). Only three of the twelve cell lines were classified as WTY, which might explain the statistical non-significance. Notably, this underrepresentation of cell lines with WTY status is consistent with our observations in established PDAC cell lines (Fig. 2a) and further supports LOY as a positively selected event, also in cell culture.

We next extended our analysis to a cohort of patient-derived organoids (PDOs) for which RNA-seq data were available^46^. Stratification of the 13 male PDOs using the rLOY signature identified 6 WTY, 2 mosaic, and 5 LOY cases (Fig. 6a), reflecting the frequency of LOY in patients. As before, the mosaic PDOs were excluded from downstream analyses for clarity reasons. When comparing treatment responses in WTY and LOY PDOs (Fig. 6b) to gemcitabine, irinotecan, and Nab-Paclitaxel (Fig. 6c-e), LOY PDOs continuously exhibited reduced sensitivity across all tested therapies (Fig. 6f). In summary, across both model cohorts, WTY alone did not appear sufficient to predict treatment response, whereas LOY emerged as a clear indicator of treatment resistance.

**Figure 6.**
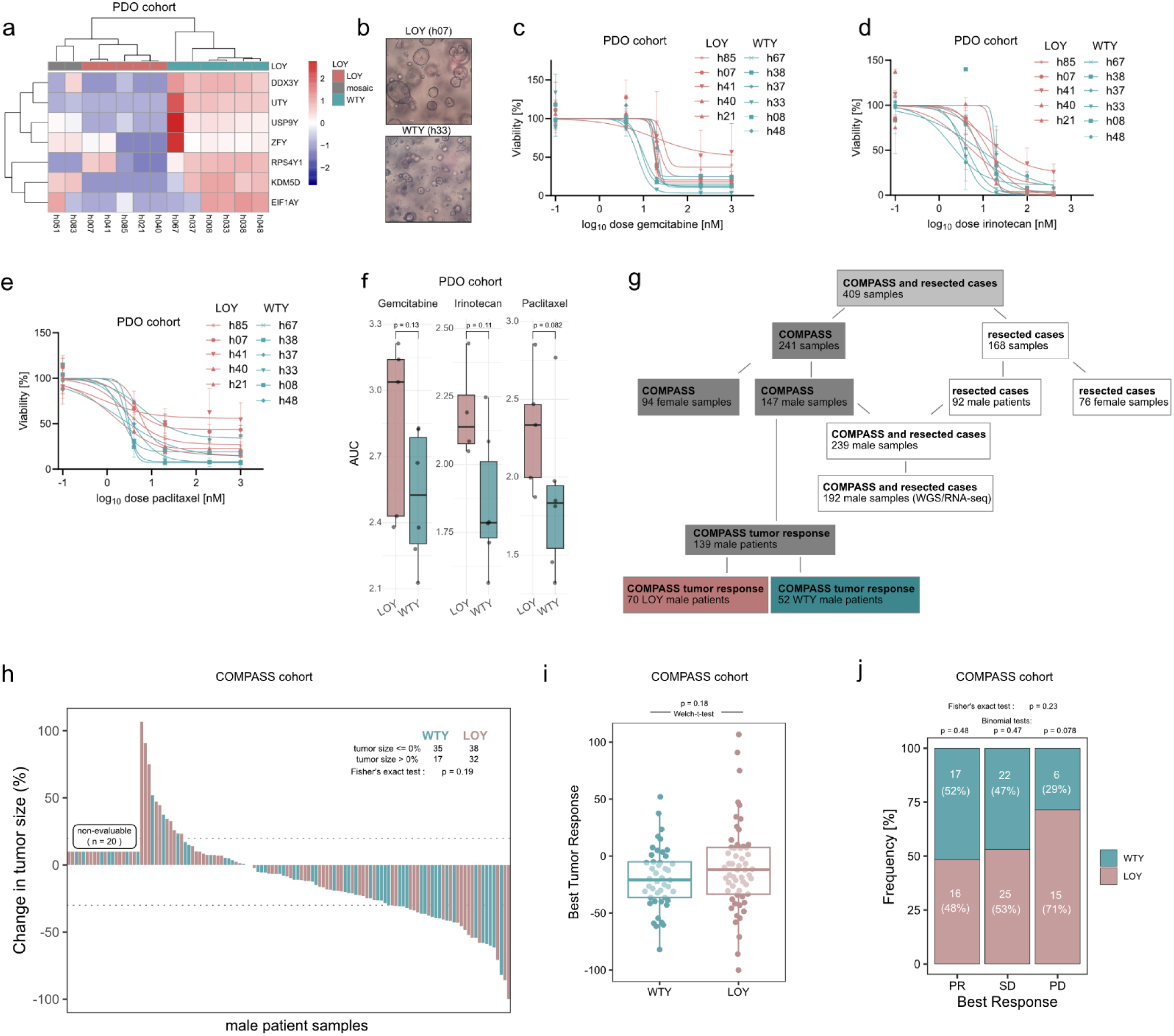
Assessment of LOY as a stratifier for treatment response. **a)** Relative expression of rLOY signature genes in 13 male patient derived organoid (PDO) lines based on RNA-Sequencing. **b)** Brightfield images of one WTY (h33) and one LOY (h07) untreated example PDO. Scale bar 100 µm. **c-e**) Viability analysis of LOY and WTY classified PDOs under treatment with irinotecan (c), gemcitabine (d), and paclitaxel (e). Data from two biological replicates are shown; for two PDOs (h08, h48), only one biological replicate was available. **f)** Comparison of AUC (area under the curve) parameters obtained from cell viability analysis in c-e. Each dot represents one PDO. Statistical significance was determined using Mann-Whitney Test. Data from two biological replicates are shown. **g)** Consort diagram of patients enrolled and included from the COMPASS and resected cases trials. 139 male patients from the COMPASS cohort were eligible for inclusion in best tumor response analysis, with WGS and RNA-Seq data as well as clinical best response data available. **h)** Waterfall plot demonstrating tumor size change according to LOY status. Tumor size change in all patients included (n = 122): This includes any chemotherapy received. Statistical significance was determined using Fisher’s exact test: p = 0.19. **i)** Comparison of best response rates in LOY versus WTY patients in the COMPASS cohort (n = 118). Statistical significance was determined using Welch-t-test, p = 0.18. **j)** Analysis of frequency of LOY and WTY patients with partial reduction (PR, p = 0.48), stable disease (SD, p = 0.47), and progressive disease (PD, p = 0.078) according to RECIST criteria. Statistical significance was determined using Fisher’s exact test and binomial tests.

Having established LOY-associated patterns across two independent patient-derived model cohorts, we next examined the primary patient cohorts. Although cohort sizes were inherently limited, reflecting the focus on male patients with complete clinical annotation and available WGS data, these datasets enabled survival and treatment-response analyses.

In the COMPASS cohort (stage III/IV, 123 male patients with corresponding survival data), overall survival (OS) did not differ by LOY status, as reflected in the Kaplan–Meier curves (Suppl. Fig. 6d, p = 0.24). However, in the resected cohort (stage I/II, 66 male patients with corresponding survival data) WTY patients demonstrated prolonged OS (Suppl. Fig. 6e, p = 0.024), suggesting that early LOY may contribute to increased tumor aggressiveness and aligning with observations in other cancer entities^12,19,47^. We next assessed treatment response in LOY versus WTY patients. Only COMPASS patients qualified for this analysis, as paired biopsy and best-response data were uniquely available for this cohort. Among 147 male patients available after strict stratification for patients with high tumor cell content, evaluable response data to standard-of-care chemotherapy (mFFX, GnP, cisplatin/gemcitabine, or gemcitabine alone) were available for 139 individuals. Subsetting on our genetic classification (Fig. 1), 52 WTY patients and 70 LOY were evaluable (Fig. 6g). Still, across all response metrics, LOY patients consistently exhibited reduced therapeutic benefit. While overall response rate (Fig. 6h, p = 0.19, Fisher’s exact test) and best tumor response (Fig. 6i, p = 0.16, Welch’s t-test) did not reach formal statistical significance, both analyses revealed a coherent trend toward poorer outcomes in LOY patients. Notably, progressive disease (PD) was enriched in the LOY group (Fig. 6j, p = 0.078, Binomial test), reinforcing the observation that LOY contributes to diminished treatment responsiveness and progression. Taken together, LOY is consistently associated with reduced sensitivity to standard-of-care chemotherapy and shows a clear correlation with progressive disease. These findings position LOY as a clinically relevant marker of treatment resistance and highlight its potential utility as an easily accessible classifier for potential treatment failure and progression in PDAC.

## DISCUSSION

Our study identifies LOY as a recurrent and functionally relevant event in PDAC. Across two independent patient cohorts, LOY was observed in ∼50% of male patients, with significant enrichment in the basal subtype and more advanced stage tumors. Mutations in *TP53* were among the most frequent alterations in both LOY and WTY, but were significantly enriched in LOY tumors, consistent with prior reports^12^. Importantly, genomic analyses revealed that LOY is independent of other large-scale genetic events like copy number alterations, structural rearrangements, *KRAS* imbalance, or genomic instability. Overall, our study identifies Y chromosome loss as the first distinct genetic event driving male pancreatic cancer progression and metastasis^24,25^. Transcriptomic profiling consistently identified both EMT and immune-related programs as the most significantly upregulated pathways in LOY tumors. Upregulation of immunoregulatory pathways aligns with prior studies linking LOY in both cancer and immune cells to altered immune surveillance and compromised tumor clearance^2,48^. This also mirrors epidemiological observations that mosaic LOY is one of the most common somatic events in hematological cells in aging men and is strongly associated with cancer, cardiovascular disease, and neurodegeneration^11,49,50^. Interrogation of single-cell RNA-seq data of male PDAC patients validated these observations in the tumor microenvironment. While there was some regulation of Y gene expression also in cells of TME, we demonstrated that LOY in malignant tumor cells primarily drives EMT. Functional interrogation using CRISPR-mediated Y chromosome knockout in PDAC cell lines demonstrated that LOY is sufficient to functionally induce conversion of epithelial cells into a mesenchymal state, highlighting that EMT signaling is the primary difference between LOY and WTY pancreatic cancer cells. This was evident at multiple levels: transcriptional activation of hallmark EMT genes, morphological changes in 2D and 3D culture, and acquisition of mesenchymal traits such as anoikis resistance and impaired colony formation. *In vivo,* LOY clones displayed reduced tumor-forming capacity in subcutaneous models, consistent with the requirement for MET during tumor establishment. Importantly, LOY cells exhibited enhanced survival in the bloodstream and increased metastatic seeding to the lung. These findings highlight the duality of EMT in promoting dissemination while impairing growth in conditions that favor epithelial-like-cells^34^.

LOY malignant cells with EMT features were enriched for signatures of the basal subtype, whereas WTY cells retained transcriptional programs of the classical subtype. While the link between (partial) EMT and the basal subtype in PDAC has been established previously^37,38^, we here identify LOY as a genetic driver of EMT and a programming towards a basal subtype in PDAC. The link between LOY and the basal, more aggressive PDAC subtype is reinforced by studies associating LOY with poorer clinical outcomes across other cancer types^19,21,51^. Interestingly, male patients present with the basal subtype of PDAC more frequently^39,52^, which can now be mechanistically explained by our finding that LOY drives the basal subtype. The mechanisms underlying the functional link between EMT and the basal subtype in female patients and how they contribute to clinical observations, remain important questions for future investigation.

Mechanistically, our study implicates epigenetic remodeling as a key mediator of LOY-induced EMT. Chromatin-modifying enzymes were consistently upregulated in LOY clones, and STRING network analyses linked these factors to EMT and ECM programs. The loss of Y-linked regulators such as *KDM5D* and *UTY* (coding for KDM6C) is particularly notable, given prior evidence that these lysine histone demethylases constrain chromatin remodeling and suppress tumor progression^23,42,43^. They regulate H3K4 and H3K27 methylation and thereby influence chromatin accessibility at lineage- and state-defining gene programs^41,44^. Here, we demonstrate that LOY induces genome-wide alterations in DNA methylation, particularly at sites that are major regulators of a mesenchymal-like cell state. Furthermore, genes defining the basal subtype, which were expressed in the YKO clones, exhibited hypomethylation, whereas those associated with the classical subtype became hypermethylated in response to LOY, further highlighting the mechanistic link between both EMT and basal subtype induction upon LOY. Thus, while the underlying epigenetic mechanisms might require more detailed investigation, LOY may act as an epigenetic switch, destabilizing transcriptional networks, and simultaneously deregulating chromatin and changing tumor cell states.

Together, these findings position LOY as a novel driver of PDAC biology in male patients, linking sex-chromosome loss to transcriptional plasticity, immune evasion, and tumor progression. Beyond PDAC, LOY has already been implicated in genitourinary cancers, lung cancer, and hematopoietic malignancies^12,22,47,53,54^, raising the possibility that Y chromosome integrity broadly constrains EMT and metastatic potential. Clinically, the high prevalence of LOY in PDAC and its association with aggressive basal tumors as well as a with reduced sensitivity to standard of care chemotherapy and progressive disease highlight its potential as a biomarker for patient stratification. This is particularly relevant for clinical translation, as FISH for the Y chromosome on an FFPE tissue slide is far more affordable and readily accessible in routine diagnostics than RNA-expression–based molecular subtyping. It also offers the added benefit of spatial resolution, enabling the distinction of LOY in tumor cells versus those in the tumor microenvironment. Moreover, the epigenetic vulnerabilities uncovered in LOY clones may provide therapeutic opportunities, particularly in targeting chromatin regulators that mediate EMT.

Across two patient cohorts, we observe a consistent trend linking loss of the Y chromosome to progressive disease and poorer clinical outcome. While statistical significance is limited, largely due to the scarcity of well-annotated LCM or single-cell RNA-seq pancreatic cancer datasets and the restriction of LOY analyses to male patients with stratification for high tumor cell content, the robustness of the effect direction underscores its potential biological relevance. Despite these constraints, our analyses in two independent clinical cohorts, 147 male patients from the COMPASS trial and 92 male patients from a resected tumor cohort, consistently point to LOY as a marker of more aggressive disease biology. Importantly, these clinical observations are reinforced by two complementary preclinical model systems, including primary patient-derived cell lines and patient-derived organoids. Together, these data provide converging evidence that LOY functions as a biologically meaningful stratifier of treatment resistance. Future studies should explore whether restoration of Y linked gene function or pharmacological modulation of epigenetic pathways can reverse LOY-induced mesenchymal states and improve therapeutic response.

Importantly, while LOY has previously been recognized as a frequent event in PDAC, our study demonstrates that complete Y chromosome loss directly drives EMT and shapes tumor behavior. This establishes LOY not merely as a correlative biomarker but as a causal determinant of PDAC progression in male patients.

## METHODS

### RESOURCE AVAILABILITY

#### Lead contact

Further information and requests for resources and reagents should be directed to the lead contact, Barbara M. Grüner. Requests for access to the COMPASS and resected cases cohorts corresponding clinical data should be directed to Faiyaz Notta. WGS and RNA-Seq data of the COMPASS and resected cases cohorts are available at https://ega-archive.org/ EGA accession number EGAS00001002543.

### EXPERIMENTAL MODEL AND SUBJECT DETAILS

#### *In vivo* animal studies

All animal experiments were performed in accordance with and approved by the Landesamt für Verbraucherschutz und Ernährung Nordrhein-Westfalen (LAVE, former LANUV, AZ 81-02.04.2019.A291 and 2024-509-GA).

Immunodeficient NOD/Scid/ᵧC (NSG) mice were originally obtained from The Jackson Laboratory via Charles River (Stock Number #005557). Animals were kept in stable groups of 2-5 animals in individually ventilated cages with standard bedding. Water and standard laboratory mouse diet were provided ad libitum. Water was filtered into autoclaved bottles. Bedding was regularly changed under a hood to preserve sterile conditions. Light/dark-rhythm was set at 12:12 hours, and room temperature and humidity were kept stable.

For all animal experiments, both male and female adult mice from the age of 8 weeks or older were equally divided and randomly assigned between all experimental groups. All animal experiments and data analyses were performed without blinding, and no power analysis was conducted to pre-determine group size. All animals were tagged with ear marks for identification prior to the experiments. No further procedures were performed besides the described experiments in this manuscript.

#### Cell lines and primary cultures

All cells were cultured under a humid atmosphere at 37 °C and 5% CO_2_. All cell culture experiments were performed in three independent biological experiments if not mentioned elsewise. Human YAPC cells (DSMZ Cat# ACC-382, RRID:CVCL_1794, male) were cultured in DMEM supplemented with 10% FBS, 1% non-essential amino acids, 50 U/ml penicillin and 50 µg/ml streptomycin. Human HupT3 (DSMZ Cat# ACC 259, RRID:CVCL_1299, male) were cultured in RPMI supplemented with 10% FBS and 50 U/ml penicillin and 50 µg/ml streptomycin. All human cell lines and derived clones have been authenticated by STR analysis. All cell lines were regularly tested to be mycoplasma negative.

### METHOD DETAILS

#### *In vivo* experiments

As stated above, all animal experiments were performed in accordance with and approved by the LAVE in the state of North-Rhein-Westphalia, Germany (Landesamt für Verbraucherschutz und Ernährung NRW).

For subcutaneous injections 0.015 x 10^6^ YAPC, HupT3 and thereof-derived YKO clones in 100 μL PBS/matrigel 1:1 were injected into each flank of NOD/Scid/ᵧC (NSG) mice. Subcutaneous tumors were measured twice weekly and experiments were terminated with a tumor size around 1000 mm^3^. Tumor weight was taken at the endpoint of the experiment.

To assess *in vivo* survival in the blood stream, 1 x 10^6^ cells in PBS were injected intravenously into the lateral tail vein of NOD/Scid/ᵧC (NSG) mice. 5 minutes after injection mice were sacrificed and analyzed for the number of cancer cells in the lung. Lung lobes were imaged from both sides using the AxioVert stereomicroscope (Zeiss). Then, two lung lobes were preserved in 4% formalin for histological analysis, the other three lung lobes were minced using scissors and digested for 0.5 hour at 37 °C in digestion media with 10% trypsin (0.25% in EDTA, Invitrogen), 10% collagenase IV (10 mg/mL in HBSS, Worthington) and 10% dispase (Corning) in HBSS w/o Ca^2+^ and Mg^2+^. Digest was quenched with L15 media (Invitrogen) containing 10% FBS and DNase (5 mg/mL in HBSS) and suspension was filtered through a 40 μm mesh, centrifuged, washed, and filtered again. Samples were analyzed for GFP expression by FACS on a Fortessa Celesta analyzer (BD Biosciences). For all samples, gating was performed on live, single cells, and one million events were acquired. No statistical method was used to predetermine sample size.

#### Histology and immunohistochemistry

Tissue samples were fixed in 4% formalin and paraffin embedded. Hematoxylin and eosin staining was performed using standard methods. Immunohistochemistry was performed using standard methods and standard antigen unmasking (1 mM citrate buffer pH 6). Primary immunoblotting antibodies were against Ki67 (Abcam Cat# ab15580, RRID:AB_443209, 1:200 dilution), and VIMENTIN (Cell Signaling Technology Cat# 5741S, RRID:AB_10695459, 1:500).

#### Viral transduction of cells

To generate YAPC and HupT3 cells labeled with GFP we used MSCV retroviral vectors generated using gag/pol-retro and VSV-G-retro packaging plasmids as previously described.^33^ For virus production, HEK293T cells were seeded into 6-well plates and individual wells were transfected at 80% confluency with plasmid vector and packaging plasmids using TransIT^®^-TKO or TransIT^®^-LT1 Transfection Reagent (Mirus). Media was changed 24 hours later. Supernatants were collected at 48 and 72 hours, pooled, centrifuged for 10 minutes at 13,200 rpm and the undiluted supernatants were each applied to 70% confluent cells. Two days after infection the cells were selected with puromycin (1 μg/mL), which effectively killed all uninfected cells. After puromycin selection, GFP expression was confirmed using flow cytometry.

#### CRISPR-Cas9 Y chromosome knockout

Cells were transfected by electrotransfer of Cas9 protein and gRNAs using a 4D-Nucleofector™ X Unit (Lonza). 2×10^5^ cells were mixed with 45 pmol of Cas9 protein (Lonza); 90 pmol of equimolar gRNA mixture and 100 pmol Alt-R Cas9 Electroporation Enhancer in Nucleofector X Solution from 4D-Nucleofector™ X Kit S (Lonza, Cat# V4XC-2032, Cat# 234005544, Cat# 234005543). Electroporation was performed using the program CM-130. Following electroporation cells were diluted into culture medium and incubated at 37°C, 5% CO_2_ in the presence of 10 µM ATM inhibitor KU-55933 for 16 h.

Equimolar gRNA mixture of targets was used at the same amounts as previously described.^29^

#### Spheroid formation assay

To initiate formation of 3D spheroids cells were seeded in triplicate (10 000 cells/well) into commercially available cell culture plates with cell-repellent surface (CELLSTAR®, 650970) in medium with drug or vehicle control treatment. Spheroids were grown for 24 h and pictures taken using Primovert microscope (Zeiss).

#### Anoikis survival assay

To assess anoikis resistance of cells and to prevent cells from adhering, 6-well plates were coated using a PolyHEMA (Sigma-Aldrich) stock solution (20 mg/ml in 95% EtOH). Cells were counted and seeded in triplicate in coated wells and also in uncoated wells as apoptosis control. After 24 h of incubation cells were stained for Annexin V (BD Biosciences Cat# 556419, RRID:AB_2665412) to monitor apoptosis using a BD FACSCelesta™ Analyzer (Becton Dickinson) as described above. For analysis, the percentage of apoptotic cells of the coated wells was normalized to the uncoated wells.

#### Colony formation assay

Cells were seeded in triplicate at 200-500 cells per well in a 6-well plate. Cells were cultivated at 37 °C up to three weeks until colonies were visible. Colonies were washed with PBS and stained for 10-30 minutes with Crystal violet. Colonies were automatically counted using Cytation5.

#### Multiplexed Y chromosome PCR

PCR was performed as previously described using the following primer sequences^31^. To visualize the correct size of PCR products, the GeneRuler DNA Ladder Mix (ThermoFisher, Cat#SM0333) was used.

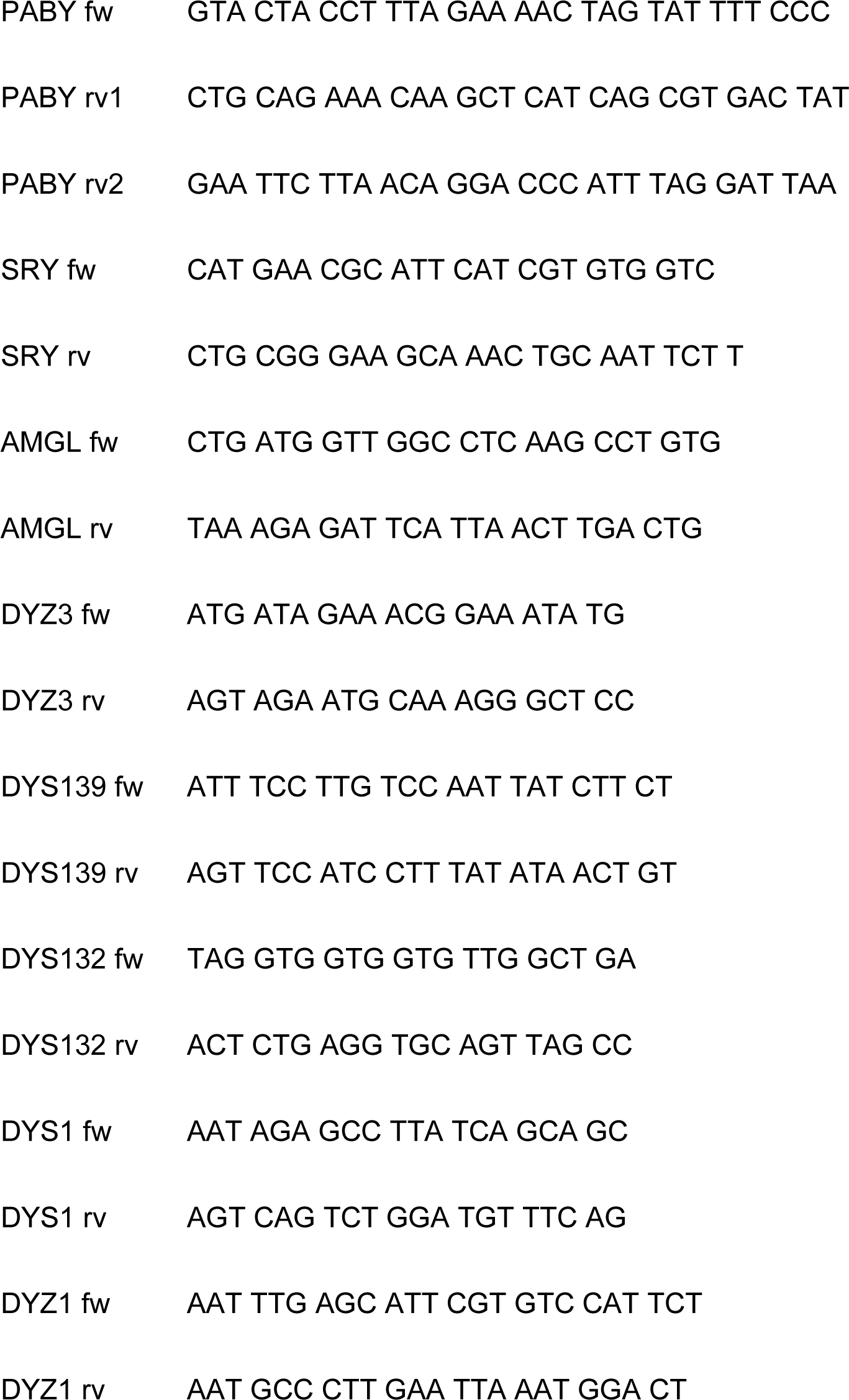

#### FISH of YAPC-YKO cell lines

Cell pellets were harvested, washed with PBS and resuspended in 4% PFA for fixation. Fluorescence in situ hybridization for Y chromosome was carried out by using the ZYTOLIGHT CEN Y/X (Zytovision, Cat#Z-2120-200) and ZytoLight CEN X/Yq12 Dual Color Probe (Zytovision, Cat#Z-2016-200) according to standard protocols.

#### Drug screening of patient derived organoids

The organoids were dissociated before plating 1000 cells in 10 μL Matrigel per well in white 96-well plates (Greiner). Cytotoxic drugs were dissolved in DMSO (concentrations were normalized to 0.25 % DMSO) and added 72 h after plating. All drugs were tested in duplicates at concentrations ranging from 1.0 × 10−7 to 1.0 × 10−3 mol/L for irinotecan (Sigma) and from 1.0 × 10−10 to 1.0 × 10−6 mol/L for gemcitabine (Sigma) and paclitaxel (Selleckchem). After 96 h of treatment, cell viability was assessed using the CellTiter-Glo 3D cell viability assay (Promega). A four-parameter log-logistic function with an upper limit equal to the mean of the DMSO values was fitted to the drug response curve and IC50s were calculated using GraphPad Prism software.

#### RNA-seq of patient derived organoids

RNA-seq was performed as previously described^46^. Sequencing libraries were prepared using the Illumina TruSeq mRNA stranded Kit following the manufacturer’s instructions. Briefly, mRNA was purified from 500 ng of total RNA using oligo(dT) beads. Then poly(A)+ RNA was fragmented to 150 bp and converted to cDNA. The cDNA fragments were then end-repaired, adenylated on the 3′ end, adapter ligated and amplified with 15 cycles of PCR. The final libraries were validated using Qubit (Invitrogen) and Tapetstation (Agilent Technologies). 2 × 100 bp paired-end sequencing was performed on the Illumina NovaSeq 6000 according to the manufacturer’s protocol. At least 54 Mio. reads per sample were generated.

#### Drug screening patient-derived cell lines

The generation of primary PDAC cells^55,56^ from PDAC specimen was approved by the ethical review board of the University Medical Center (11/5/17). Cells used in this study were derived from two independent sources: (i) patient-derived xenografts (PDX), which were further expanded as cell-derived xenografts (CDX), and (ii) patient-derived organoids (PDO), which were subsequently differentiated into organoid-derived cells (ODC). The drug screening was performed using a standardized workflow, simply as follows. A compound library of approximately 200 drugs was pre-arrayed with gradient dilutions ranging from 10 mM to 13.7 μM and stored at −20 °C. Cells were seeded in duplicates in white-bottom 96-well plates 24 hours before treatment to allow attachment. Drug dispensing was carried out using a manual pin tool (V&P Scientific), delivering compounds to achieve final assay concentrations from 10 μM to 13.7 nM. After 72 hours of incubation, cell viability was quantified using the CellTiter-Glo assay (Promega) and measured with a microplate reader. For this study, only responses to standard chemotherapies were reported. Relative viability, growth-rate–based dose–response curves, and summary metrics including AUC were calculated using the GRmetrics R package.

#### COMPASS and resected PDAC patient cohorts

Whole-genome and transcriptome libraries were previously sequenced, as described^27^. To estimate copy number profiles, B-allele frequencies were estimated using germline single nucleotide variant calls derived using GATK’s v. 4.1.2 haplotypecaller^57^. Read depth was segmented using HMMCopy 0.1.1 and integrated with B-allele frequency to derive copy number alterations, tumor content and ploidy using Celluloid v0.11.7^58^. All celluloid solutions were manually reviewed.

Germline sex chromosome complement was inferred using the ratio of reads mapped to the X chromosome over those mapped to the Y chromosome in a buffy coat (normal) sample. Individual sex chromosome class (XX versus XY) was called by binning a Bayesian mixture model fitted on cohort-level X/Y ratios with MClust v.6.1.1. To call Y chromosome copies, Y chromosome segments were bound between positions 11,659,001 and 26,518,001 of hg38; remaining Y regions were excluded due to poor assembly and high repetitive content^59^. LOY was defined using a similar procedure as germline sex chromosome calls: for each individual, we calculated the median absolute copy number of Y segments estimated by *Celluloid* and fit a mixture model to the cohort distribution, preset to four latent distributions (0, 0-1, 1, 1+). Samples in the bin defined by median Y copy number between 0 and 1 had their complete Y copy number profile manually reviewed and were found to have partial Y deletions and were removed from analysis. Samples with more than one Y chromosome were mostly whole genome duplicated: Y copy gain without whole genome duplication occurred in <2% (*n* = 4) of cases. Difference in counts of structural variants was used as a measure of genome instability and compared between groups using a Wilcoxon rank sum test. Associations between mutation status or KRAS imbalance and LOY were assessed using Fisher’s exact test with multiple testing correction.

RNA reads were mapped per transcript to ensembl version 100 of hg38.p14 using star 2.7.4a, to a median of 45 million unique fragments per sample^60^. Reads were tallied per gene and normalized by coverage and gene length to produce transcript per million (TPM) values using *stringtie* v2.0.6. An LOY classifier based on the seven genes with discriminative power identified by (Qi et al. 2023) was re-trained for PDAC^12^. To do this, we trained a binomial generalized linear model on a balanced subset of 60 WGS-defined LOY samples and 60 WTY samples, and tested precision and recall on the remaining samples in the cohort. For differential gene expression analysis, reads were tallied per gene using *htseq*v2.0.4^61^ and subset to protein coding genes longer than 500 bp on chromosomes 1-22 before normalization by DeSeq2 v1.48.1. Differential gene expression analysis was performed with DeSeq2 and gene set enrichment analysis was performed with *GSEA* in clusterProfiler v4.16.0 using the Hallmarks set.

#### Whole-genome sequencing of YKO cell lines with CeGaT

Tumor DNA from three PDAC cell lines (YAPC, YAPC-YKO-1, YAPC-YKO-3) (600 ng each) was submitted to CeGaT GmbH (Tübingen, Germany) for whole-genome sequencing (WGS). DNA quality was assessed via fluorescence-based quantification, and all samples passed quality control (concentrations: 20.3–22.3 ng/μl). Library preparation was performed using the TruSeq DNA Nano kit (Illumina) with 100 ng input per sample. Sequencing was conducted on the NovaSeq X Plus platform, yielding Q30 values of 95.23%.

Sequencing reads were demultiplexed using Illumina bcl2fastq (v2.20), and adapters were trimmed with Skewer (v0.2.2). Reads were mapped to the hg19 reference genome using the Illumina DRAGEN Bio-IT Platform (v4.2.4); duplicates were marked, and no additional quality trimming was applied. Median insert sizes ranged from 381 to 394 bp, and average genome coverage was 30.6–31.3×.

Variant calling was performed using DRAGEN default parameters. Small variants (SNVs and indels), structural variants (SVs), and copy number variations (CNVs) were identified, including mitochondrial CNVs (chrM), which were not interpreted due to cell-type variability. CNV calling was conducted in self-normalization mode. Annotation of variants was performed using Nirvana, with outputs provided in VCF, TSV, and JSON formats.

Quality metrics were compiled using MultiQC (v1.22.2), and FASTQ file quality was assessed via FastQC. Visualization and summary plots were generated in R (v4.0.4) using ggplot2. All data files are available upon request.

#### Whole-genome sequencing CNV analysis of YKO cell lines

Paired-end WGS reads were aligned to the human genome (GRCh38.p14) using BWA-MEM (v0.7.17) with default parameters. Resulting SAM files were converted to BAM format, sorted, and indexed using Samtools (v1.15).

The human genome was partitioned into consecutive 20 kb bins using bedtools makewindows, generating a genome-wide bin file. Read coverage for each bin was computed with bedtools coverage for all WGS samples. GC content for each bin was calculated using bedtools nuc.

Bin-level coverage was normalized for GC content using a LOESS regression, where normalized read depth (RD) was defined as the ratio of observed coverage to predicted coverage based on GC content. Log_2_ copy number ratios were then calculated as:

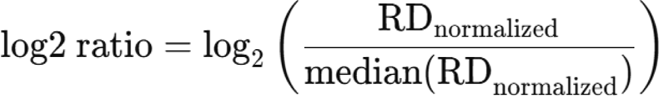

Bins with infinite log_2_ ratios were set to NA.

Normalized log_2_ ratios were visualized using pheatmap (R package v1.0.12). Samples were displayed as rows and genomic bins as columns.

#### RNA-Seq analysis of YKO cell lines by Singleron

Biological replicates of YAPC parental and YAPC-YKO-1 and YAPC-YKO-3 were harvested and cell pellets were stored at -80 °C. Snap frozen tumor cell pellets were shipped to Singleron Technologies GmbH (Cologne, Germany) for RNA isolation and bulk transcriptome sequencing service. Total RNA was isolated using the GeneMATRIX Universal RNA Purification Kit (cat#E3598-02, Roboklon). Subsequently, RNA quantity and integrity were assessed using Qubit^TM^ RNA High Sensitivity (HS) Assay Kits (cat#Q10211, Invitrogen^TM^) and High Sensitivity RNA ScreenTape (cat#5067-5579, Agilent). Purified RNA was processed according to the instructions of the AccuraCode^®^ RNA-Seq kit V2 (cat# 10710174, Singleron Biotechnologies GmbH). Briefly, polyadenylated mRNA from different samples was captured using paramagnetic beads. During reverse transcription, the mRNA was converted into cDNA and labeled with sample-specific barcodes and unique molecular identifiers (UMI). The labeled cDNA was pooled, amplified and used to construct sequencing libraries, which were sequenced on Illumina NovaSeq X using a paired-end 150 bp sequencing strategy by GENEWIZ (Leipzig, Germany).

Raw reads were processed to gene expression matrices using CeleSCOPE v2.7.4 (multi_bulk_rna workflow with customized chemistry; www.github.com/singleron-RD/CeleScope); Singleron Biotechnologies GmbH). R2 FASTQ files containing transcript sequences were demultiplexed using well-specific barcodes extracted from the corresponding R1 FASTQ files. Reads were then aligned to the *Homo sapiens* GRCh38 Ensemble 99 reference genome.

#### Single cell RNA-seq analysis

Publicly available single cell RNA sequencing data were obtained from Werba et al^35^. The cell ranger-processed output files were downloaded from the GEO repository (GSE205013), data for male patients was extracted and analyzed following the workflow described in the study, using the Seurat package (v4). Cells were filtered according to the thresholds reported in the publication: >500 detectable genes, >1,500 unique molecular identifiers, <15% of transcripts coming from mitochondrial genes, and >1% of transcripts representing erythroid genes (*HBA1, HBA2, HBB, HBM, ALAS2*). Expression counts were normalized by library size, scaled to 10,000 counts per cell, and subsequently log-transformed. Doublets were identified and removed using DoubletFinder (v2.0.6). Seurat anchor-based integration as well as dimensionality reduction of the scaled, integrated data matrix were performed following the workflow from the original publication. The main cell clusters were annotated based on marker genes described in the paper. Identification of malignant cells was performed using copyKAT (v1.1.0)^36^, with all non-epithelial cells used as reference. To account for patient-specific batch effects in male malignant cells, harmony integration prior to downstream analysis was performed. Stratification of male cells/patients into LOY or WT groups was based on expression of Y chromosome-specific genes: *DDX3Y, EIF1AY, KDM5D, RPS4Y1, USP9Y, UTY, ZFY*.

#### Pathway and network analysis

Pathway enrichment analysis of all datasets was performed using Metascape, and protein– protein interaction network analysis was conducted with STRING (Version 12.0, https://string-db.org/). For pathway analysis, only genes exhibiting a minimum absolute log_2_ fold change of 0.25 and an adjusted p-value < 0.05 were included. Visualization of the Metascape output was performed using R/RStudio.

#### Quantitative real-time PCR (qRT-PCR)

Cells were harvested and RNA was isolated using the RNeasy Mini Kit from Qiagen (Cat# 74104) and reversely transcribed as per the manufacturer’s protocols using High-Capacity cDNA Reverse Transcription Kit from ThermoFisher (Cat# 4368814).

qRT-PCR was performed using LightCycler® 480 SYBR Green I Master Kit (Roche) according to standard protocols and manufacturer’s instructions. Analysis was performed in triplicate and relative expression was calculated with the ΔΔCT method. *GAPDH* expression was used for normalization.

Primer sequences:

SNAI2: fw CGAACTGGACACACATACAGTG, rv CTGAGGATCTCTGGTTGTGGT

GAPDH: fw CATCAAGAAGGTGGTGAAGC, rv GAGCTTGACAAAGTGGTCGT

VIMENTIN: fw AGTCCACTGAGTACCGGAGAC, rv CATTTCACGCATCTGGCGTTC

#### EPIC Array and analysis

DNA methylation profiling was performed using Illumina Infinium Methylation EPIC array data comparing YAPC-YKO clones 1 and 3 (combined) versus YAPC-parental cells. Raw IDAT files were processed and analyzed using RnBeads, a comprehensive bioinformatics pipeline specifically designed for the analysis of Illumina methylation arrays. RnBeads performs quality control, normalization, probe filtering, statistical testing, and genomic annotation in a fully integrated workflow, enabling robust identification of differentially methylated CpG sites and regions. In brief, low-quality probes, probes overlapping single nucleotide polymorphisms (SNPs), and probes mapping to multiple genomic locations were excluded prior to downstream analyses. Differential DNA methylation analysis was carried out using RnBeads’ statistical framework, comparing YAPC-YKO clones 1 and 3 against parental YAPC cells. CpG sites were considered statistically significant if they fulfilled both of the following criteria: (i) an absolute mean methylation difference (Δβ) greater than 0.20 between groups, and (ii) a p-value < 0.05. Using these thresholds, a total of 36,869 CpG sites were identified as significantly differentially methylated between YAPC-YKO and parental cells. To characterize the functional genomic context of the differentially methylated CpGs, all significant CpG sites were annotated using ChromHMM chromatin state segmentation, which integrates multiple histone modification marks to define distinct regulatory states across the genome. This analysis classified CpG sites into 16 different chromatin states, including promoters, enhancers, transcribed regions, and repressed or heterochromatic regions. Hypermethylation indicates increased methylation in YAPC-YKO cells, whereas hypomethylation indicates decreased methylation relative to controls.

#### Statistical analysis

Graphs were generated using GraphPad Prism or R/RStudio, and statistical analyses were predominantly performed with non-parametric tests, as specified in the corresponding figure legends.

## RESOURCES TABLE and DATA AVAILABILITY

Rstudio and R 4.5.1; Seurat package (v4); DoubletFinder (v2.0.6); copyKAT (v1.1.0); CeleSCOPE v2.7.4; BWA-MEM (v0.7.17) ; Samtools (v1.15); bedtools nuc; Illumina bcl2fastq (v2.20), Skewer (v0.2.2); Illumina DRAGEN Bio-IT Platform (v4.2.4); GSEA in clusterProfiler v4.16.0; DeSeq2 v1.48.1; stringtie v2.0.6; htseqv2.0.4; GATK’s v. 4.1.2 haplotypecaller; HMMCopy 0.1.1; Celluloid v0.11.7; MClust v.6.1.1

RESOURCES TABLE and DATA AVAILABILITY
| REAGENT or RESOURCE | SOURCE | IDENTIFIER |
| --- | --- | --- |
| **Antibodies** |  |  |
| Rabbit polyclonal anti Ki67 | Abcam | Cat# ab15580, RRID:AB_443209 |
| Rabbit monoclonal anti Vimentin (D21H3) | Cell Signaling Technology | Cat# 5741, RRID:AB_10695459 |
| **Deposited data** |  |  |
| Werba et al. | 35 |  |
| RNA-seq YAPC-YKO clones | This paper | GEO record number: GSE319262 |
| COMPASS and resected PDAC cases | 27 |  |
| EPIC-array YAPC-YKO clones | This paper | GEO record number: GSE318463 |
| WGS-seq | This paper | GEO record number: GSE319263 |
| **Experimental models: Cell lines** |  |  |
| Human YAPC cells | DSMZ | ACC-382 |
| Human HupT3 cells | DSMZ | ACC-259 |
| **Experimental models: Organisms/strains** |  |  |
| Immunodeficient NOD/Scid/γC (NSG) mice | The Jackson Laboratory via Charles River | #005557 |
| **Oligonucleotides** |  |  |
| qPCR Primers, see methods section | Eurofins | N/A |
| **Recombinant DNA** |  |  |
| MSCV-GFP; PGK Puro | Addgene | #68486 |
| VSV-G retro | Addgene | #8454 |
| gag/pol retro | Addgene | #14887 |
| **Software and algorithms** |  |  |
| Graphpad Prism | N/A | https://www.graphpad.com/company/ |
|  |  | Version 10 |
| STRING | 62 | https://string-db.org/<br>Version 12.0 |
| metascape | 63 |  |

## ACKNOWLEDGMENTS

We thank Eva Hahn and Tabea Hugo for excellent technical assistance. We thank the members of the Grüner, Grünwald, Notta and Schramm labs for helpful comments and discussions. We acknowledge support by the Westdeutsche Biobank Essen (WBE, University Hospital Essen, University of Duisburg-Essen, Essen, Germany. We also thank the central animal facility of the University Hospital Essen (ZTL) for their support. This work was supported from the BMFTR SATURN^3^ joint project funded by the National Decade Against Cancer (01KD2206A), and the Deutsche Forschungsgemeinschaft (DFG, German Research Foundation) with an Emmy Noether Award (GR4575/1-2) both to B.M.G, by the Sander Stiftung, grant no. 2020.166.2 (to A.S. and B.M.G.), and by the Else Kröner Fresenius Stiftung (2025_EKEA.228) to M.D. F.R. acknowledges funding by the Melanoma Research Alliance and the Wolfgang & Gertrud Boettcher Foundation. R.F.S is a Professor at the Cancer Research Center Cologne Essen (CCCE) funded by the Ministry of Culture and Science of the State of North Rhine-Westphalia. This work was partially funded by the German Ministry for Education and Research as BIFOLD - Berlin Institute for the Foundations of Learning and Data (ref. 01IS18025A and ref 01IS18037A) and by the Bruno and Helene Jöster Foundation as “CLONETRAC - Tracking the clonal dynamics of cancer through treatment at the single-cell level” both to R.F.S. E.H. and G.S. acknowledge funding by DFG - CRU5002 and RTG2978. N.D. was supported by start-up-funding provided by the DFG-funded Clinical Research Unit (CRU) 5002. G.S. and B.M.G acknowledge funding by the German Cancer Aid (DKH – Deutsche Krebshilfe) DFEAT-PDAC (P02). J.P.N. declares grants from Heidelberger Stiftung Chirurgie, Dietmar Hopp Stiftung GmbH, Bundesministerium für Bildung und Forschung, and Stiftung Deutsche Krebshilfe (DKH). J.T.S. acknowledges funding by the German Cancer Consortium (DKTK) and by the German Federal Ministry of Research, Technology and Space (BMFTR; 01KD2206A/SATURN3).

## ETHICS DECLARATIONS

### Competing intrests

J.P.N. declares the patent 52378-704.601 PCT 23 May 2025, Combination of Irinotecan and AP-001 for treating cancer; and a consulting advisory role for BioNTech SE.

J.T.S. receives honoraria as consultant or for continuing medical education presentations from AstraZeneca, Bayer, Boehringer Ingelheim, Bristol-Myers Squibb, Immunocore, MSD Sharp Dohme, Novartis, Roche/Genentech, and Servier. His institution receives research funding from Abalos Therapeutics, AstraZeneca, Boehringer Ingelheim, Bristol-Myers Squibb, Eisbach Bio, Oncolytics Biotech and Roche/Genentech; he holds ownership in FAPI Holding (< 3%). All other authors declare no competing interests.

## AUTHOR CONTRIBUTIONS

Conceptualization: M.D., B.M.G.

Methodology: M.D., F.E.G.B., B.T.G., B.M.G.

Formal analysis: M.D., F.E.G.B., A.K.B., P.Z., B.O-C., N.D., L.G., M.M.C., K.S., K.G., F.O., T. J. Y. K., M.L-P.

Investigation: M.D., B.O-C., A.P., S.U., J.Z., J.M.H., J.J., N.M., H-U.S., T.P.

Visualization: M.D., F.E.G.B., A.K.B., P.Z., B.O-C., B.M.G.

Validation: M.D., B.T.G., B.M.G.

Resources and concepts: R.F.S., E.H., G.S., K.St., J.T.S., J.P.N., T.P., A.S., C.P., F.R., F.N., B.T.G.

Writing – original draft: M.D., B.M.G. Writing – review & editing: all authors Funding acquisition: B.M.G.

