## Supplementary Figures S1-S6 for "Loss of the Y chromosome drives basal identity and metastatic potential via epigenetic reprogramming in pancreatic cancer" (from another version of this preprint: version 1, 2026-10-06)

### Suppl. Figure 1

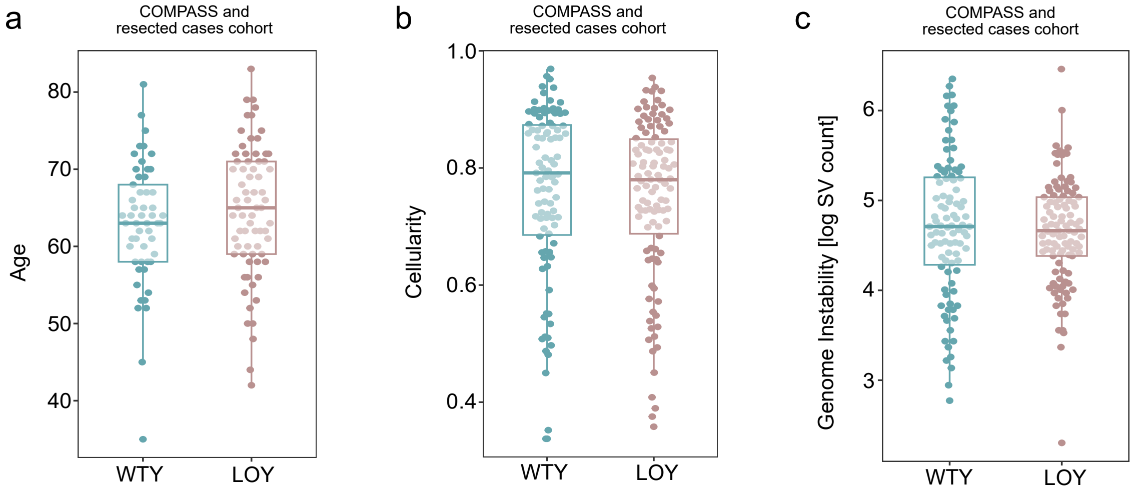

**Figure S1. Genomic consequences of Y chromosome loss (LOY) in male PDAC patients**

**a)** Distribution of age in LOY and WTY patients from the COMPASS and resected PDAC cases cohort (n = 123; p = 0.274, Wilcox rank sum test).

**b)** Distribution of cellularity (percentage of tumor cells in analyzed sample) in LOY and WTY patients from the COMPASS and resected PDAC cases cohort for whom this data was available (n = 201; p = 0.43, Wilcox rank sum test).

**c)** Analysis of genomic instability based on structural variants in LOY and WTY patients from the COMPASS and resected PDAC cases cohort for whom this data was available (n = 201; p = 0.52, Wilcox rank sum test).

### Suppl. Figure 2

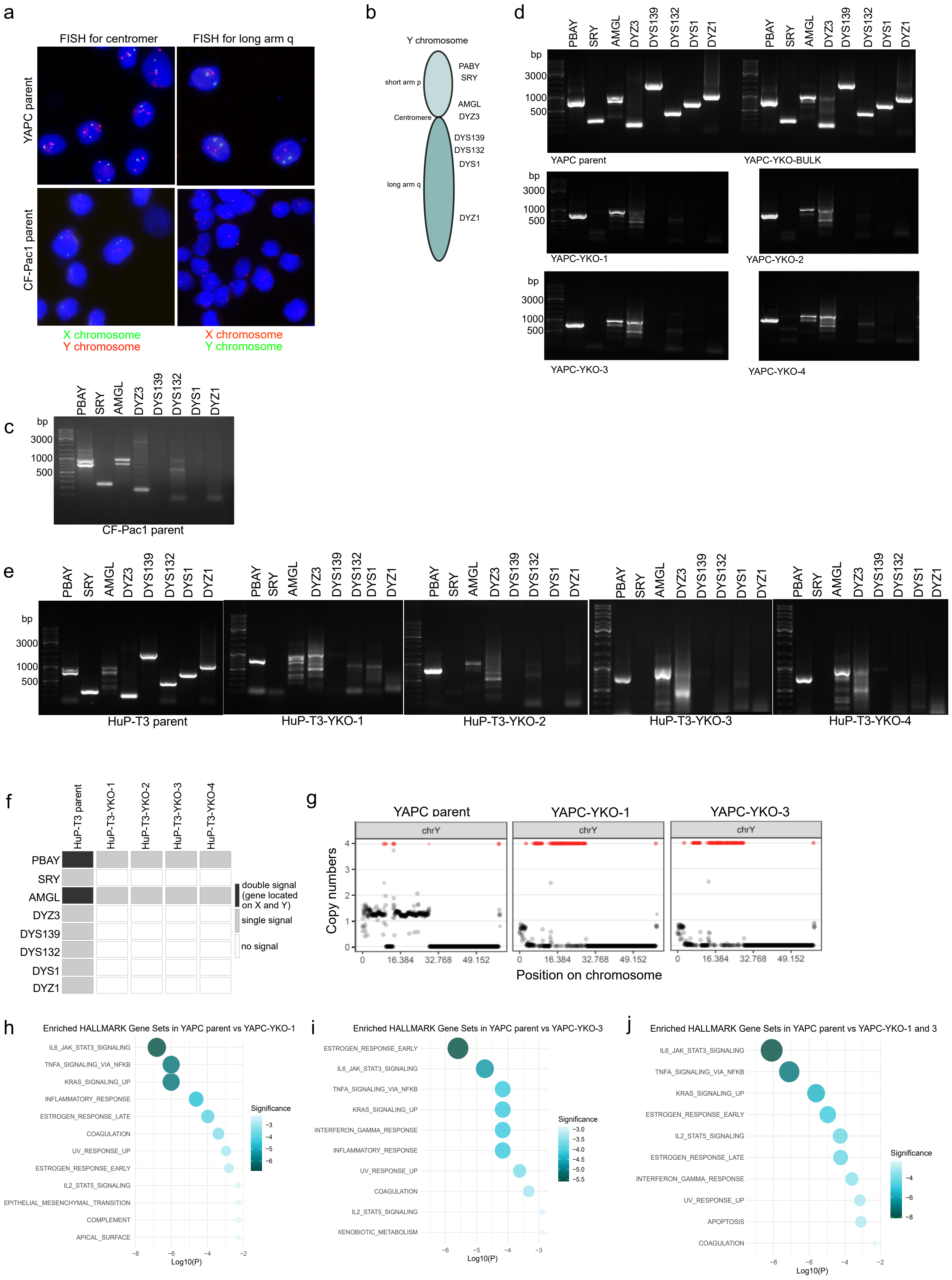

**Figure S2: Genomic and transcriptomic characterization of functional CRISPR-CAS-9-Knockout of the Y chromosome in PDAC cell lines**

**a)** Fluorescence *in situ* hybridization (FISH) analysis of the centromer (red, left panel) or the long arm q (green, right panel) of the Y chromosome in YAPC (upper panel) or CF-Pac1 (lower panel) cells. The X chromosome centromere is displayed as internal control, using dual color probes.

**b)** Schematic of the Y chromosome and the location of the respective genes amplified for the Y chromosome PCR depicted in (c-e).

**c-e)** Y chromosome PCR in CF-Pac1 (c), YAPC parent and YAPC-YKO clones (d), and Hup-T3-parent and Hup-T3-YKO clones (e).

**f)** Semi-quantitative analysis of Y chromosome multiplex PCR. Table summarizing signal detection for nine Y-linked genes across Hup-T3-YKO clones.

**g)** Whole genome analysis of the Y chromosome in YAPC-parent and YAPC-YKO-1 and YAPC-YKO-3 clones depicting relative gene coverage.

**h-j)** Pathway enrichment analysis of RNA-seq data comparing YAPC-YKO-1 (h) and YAPC-YKO-3 clones (i) alone, or both clones' overlapping genes combined (j) compared to YAPC-parent cells. Dot plot shows significantly downregulated HALLMARK pathways from genes downregulated in YKO clones ($log_2$ fold change > 0.75, adjusted $p < 0.05$).

### Suppl. Figure 3

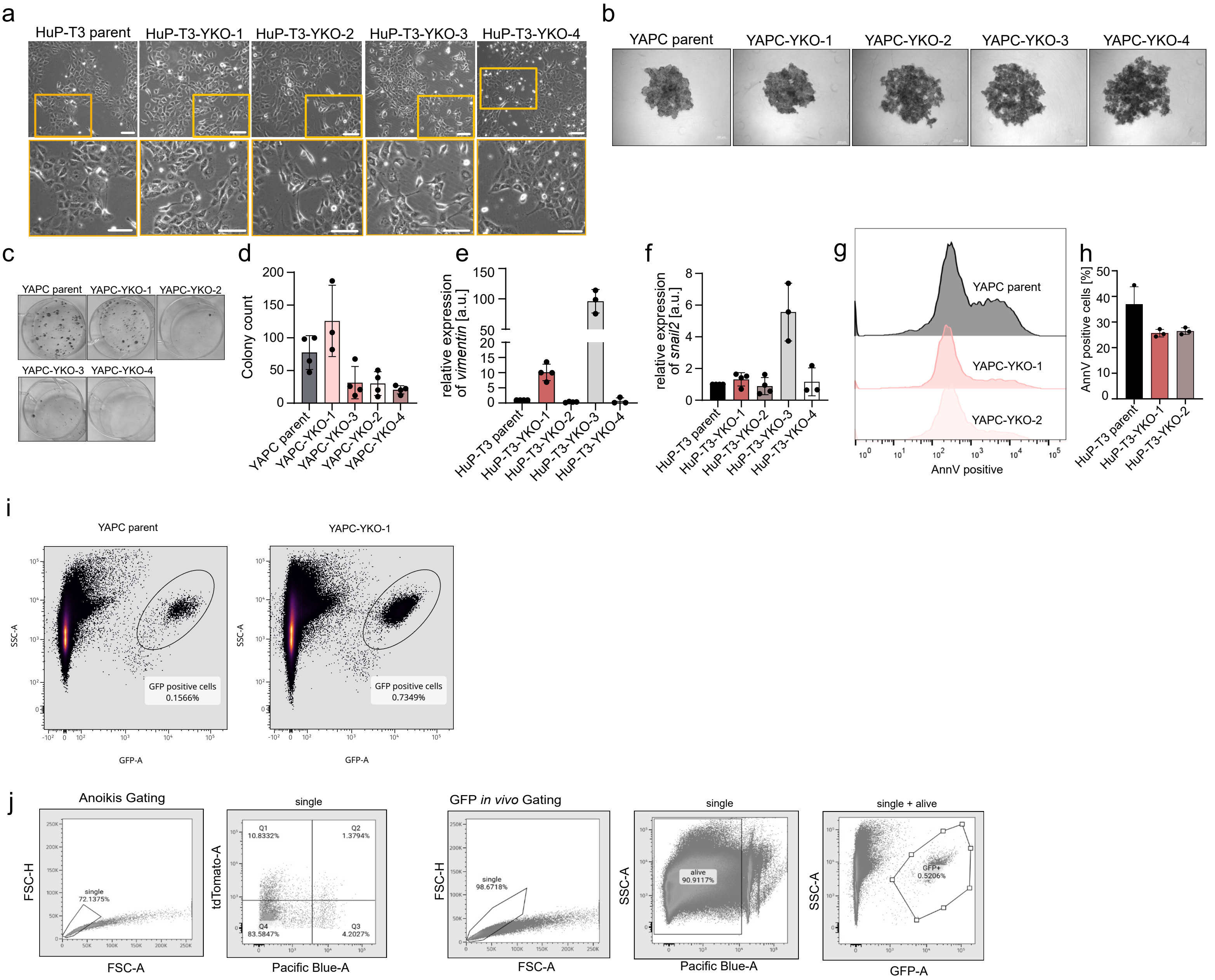

**Figure S3: Functional *in vitro* and *in vivo* analysis of YKO cell line clones**

**a)** Representative 2D morphology images of HuP-T3-parent cells and derived HuP-T3-YKO clones. Magnified regions highlighted by yellow boxes. Scale bars represent 100 µM.

**b)** Representative 3D morphology images of YAPC-parent cells and YAPC-YKO clones. Scale bars represent 200 µM.

**c,d)** Colony formation assay of YAPC-parent cells and YAPC-YKO clones. Representative images (c) and quantification of three independent experiments (d) are depicted. Statistical significance was determined using the Kruskal–Wallis test; non-significant comparisons are not indicated.

**e,f)** Quantitative RT-PCR analysis of mesenchymal markers *Vimentin* (e) and *SNAIL2* (f), normalized to *GAPDH* in Hup-T3-parent versus Hup-T3-YKO clones. Quantification of three independent experiments is depicted. Statistical significance was determined using the Kruskal–Wallis test; non-significant comparisons are not indicated.

**g)** Representative flow cytometry histogram of cells undergoing anoikis in YAPC-parent and YKO clones.

**h)** Quantification of cells undergoing anoikis in Hup-T3-parent and Hup-T3-YKO clones. One representative out of three independent experiments is depicted. Statistical significance was determined using the Kruskal–Wallis test; non-significant comparisons are not indicated.

**i)** Representative flow cytometry analysis of GFP-positive cells in lungs of mice injected intravenously with YAPC-parent or YAPC-YKO-1.

**j)** Gating strategy for anoikis assay (upper panel) and assessing GFP-positive cells in lungs injected intravenously with GFP-positive cells (lower panel).

### Suppl. Figure 4

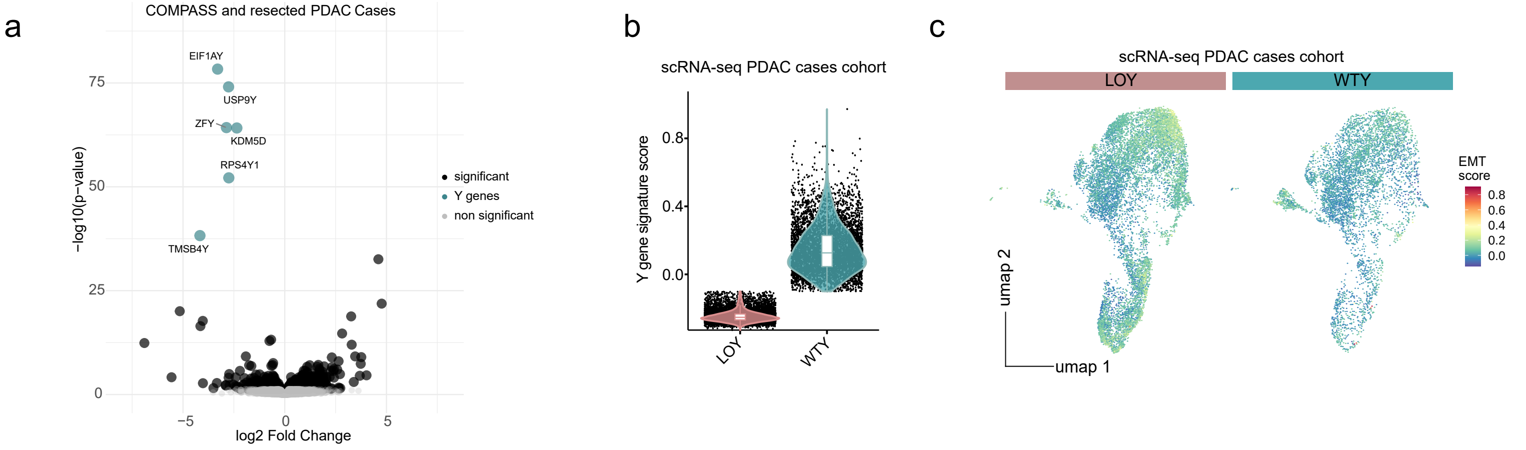


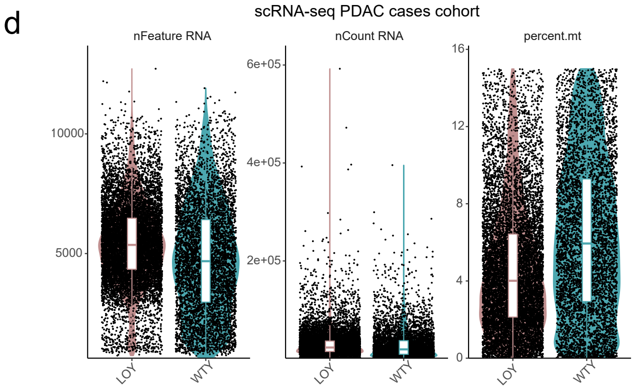

**Figure S4: Transcriptomic profiling of LOY in two independent male PDAC patient cohorts**

**a)** Differential gene expression analysis in LOY versus WTY patient samples in the COMPASS and resected PDAC patient cohort.

**b)** Y-gene signature score expression in WTY versus LOY epithelial malignant cells stratified by CopyKat in the Werba et al. single cell dataset.

**c)** UMAP of the EMT hallmark gene set in WTY versus LOY epithelial malignant cells stratified by CopyKat in the Werba et al. single cell dataset.

**d-f)** Distribution of gene number (d), UMI counts (e), or mitochondrial gene content (f) in WTY versus LOY epithelial malignant cells stratified by CopyKat in the Werba et al. single cell dataset.

### Suppl. Figure 5

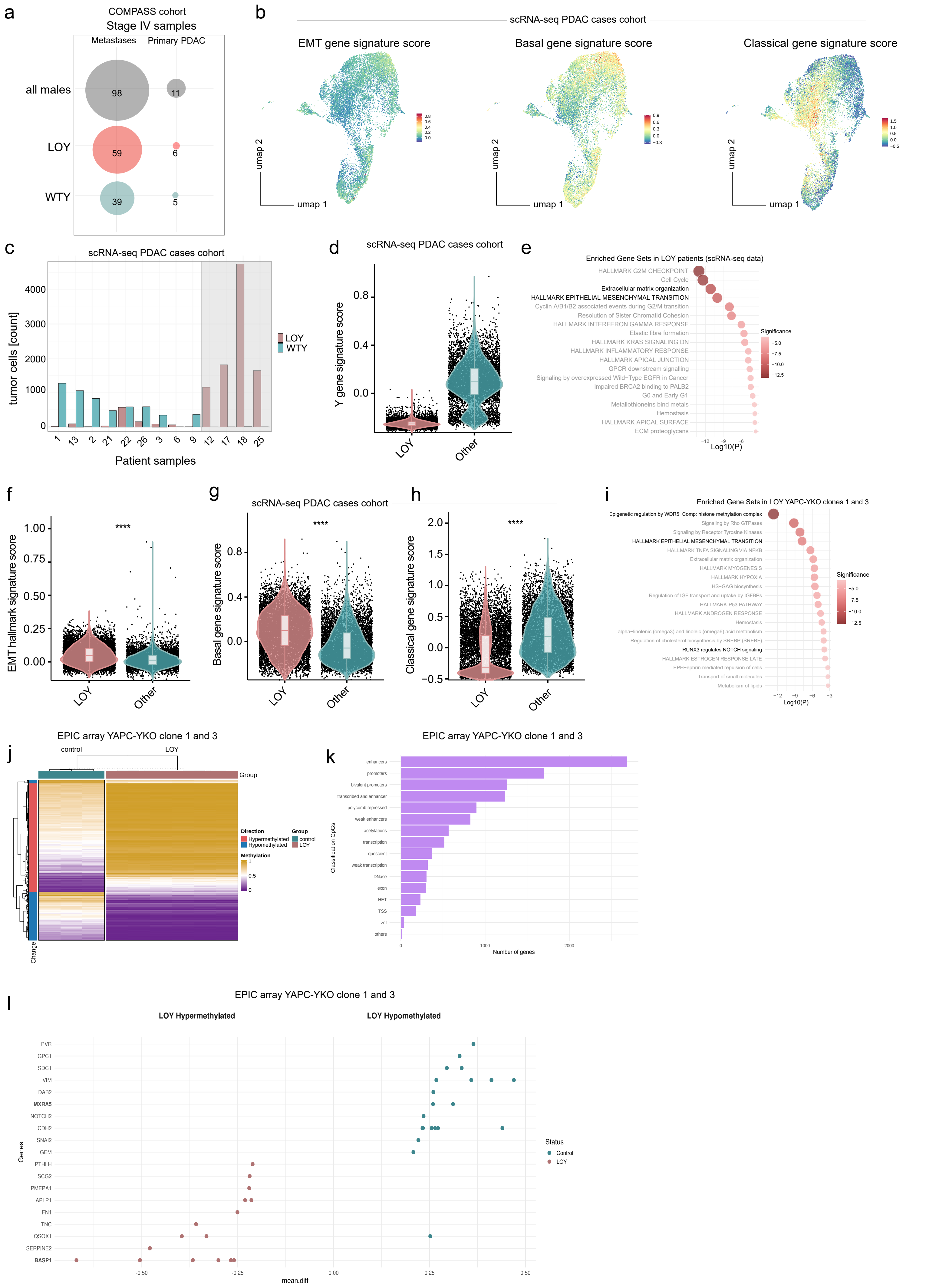

**Figure S5: Functional connection of LOY with the basal subtype via epigenetic regulation**

**a)** Distribution of LOY and WTY cases in stage IV patients from primary tumors or metastasis in the COMPASS and resected PDAC cases patient cohort.

**b)** UMAP of the EMT gene signature, Moffitt.mod basal signature and Moffitt.mod classical gene signature of epithelial malignant cells stratified by CopyKat in the Werba et al. single cell dataset.

**c)** Cell count of epithelial malignant cells per patient stratified for LOY or WTY according to the Y gene signature score in the Werba et al. single cell dataset.

**d)** Violin plot comparing Y-gene signature scores (rLOY) in epithelial malignant cells between LOY and "other" patients (stratified in c) in the Werba et al. single cell dataset.

**e)** Gene set enrichment analysis in epithelial malignant cells between LOY and "other" patients (stratified in c) in the Werba et al. single cell dataset. Dot plot shows significantly upregulated HALLMARK and Reactome pathways from genes upregulated in LOY patients ($\log_2$ fold change >0.75, adjusted $p < 0.05$).

**f-h)** Violin plots compare EMT hallmark (f), Moffitt.mod basal (g), and Moffitt.mod classical (h) gene signature scores between LOY and "other" stratified patients (c) in epithelial malignant cells in the Werba et al. single cell dataset. Statistical significance was determined using Wilcoxon Test, **** $p < 0.0001$.

**i)** Gene set enrichment analysis of hallmark and Reactome pathway signatures in both YAPC-YKO-1 and YAPC-YKO-3 clones versus YAPC-parent. Dot plot shows significantly upregulated HALLMARK and Reactome pathways from genes upregulated in both YAPC-YKO clones ($\log_2$ fold change < –0.75, adjusted $p < 0.05$).

**j-l)** EPIC Array analysis comparing combined YAPC-YKO-1 and YAPC-YKO-3 clones to YAPC-parent controls. j, Heatmap of DNA methylation levels in LOY (YAPC-YKO-1 and YAPC-YKO-3 clones) and control samples. 36,896 CpGs were filtered using mean.diff > 0.2 and diffmeth.p.val < 0.05; from these, the 500 most hypermethylated and 500 most hypomethylated sites were selected to visualize the strongest LOY-associated changes. k,

ChromHMM classification of differentially methylated CpGs across 16 chromatin states, illustrating their functional annotation. I, Nineteen overlapping EMT genes identified through CpG annotation; MXRA5 and BASP1 are highlighted as promoter-associated.

### Suppl. Figure 6

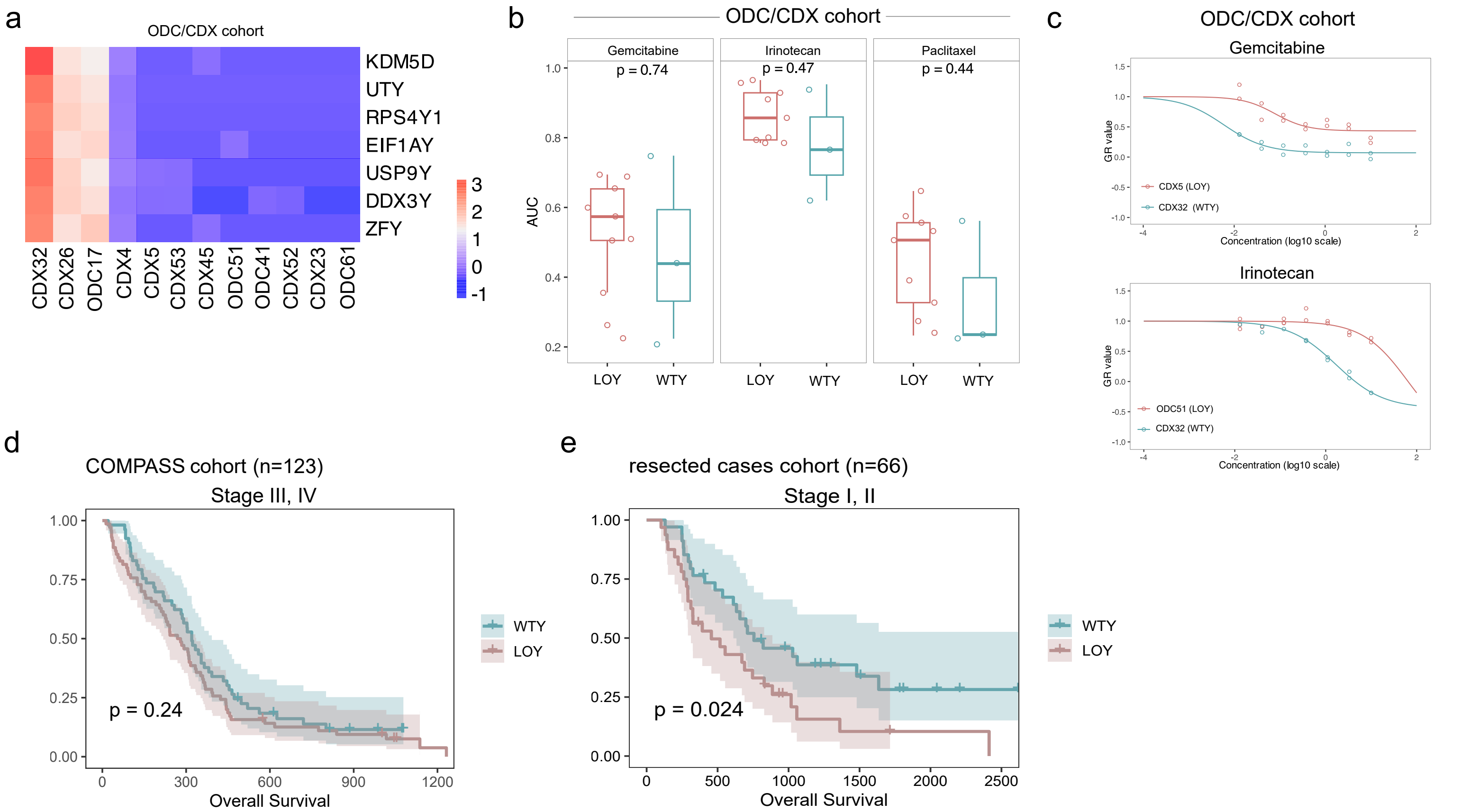

**Figure S6. Assessment of LOY as a stratifier for treatment response.**

**a**) Relative expression of rLOY signature genes in 12 male primary patient derived cell lines based on RNA-Sequencing. Cells were derived from two sources: PDX-derived CDX and PDO-derived ODC.

**b,c)** Viability analysis of LOY and WTY classified primary patient derived cell lines under treatment with irinotecan, gemcitabine, and paclitaxel. Comparison of AUC (area under the curve) parameters (b) obtained from cell viability analysis exemplified in (c). Each dot represents one primary cell line. Statistical significance was determined using Mann-Whitney Test. Data from one biological replicate is shown.

**d, e**) Overall survival (OS) analysis in the COMPASS cohort (d, n = 123 eligible male patients, p = 0.24), and the resected cases cohort (e, n = 66 eligible male patients, p = 0.024). 66 male patients from the resected cases cohort and 107 male patients from the COMPASS cohort were eligible for overall survival analysis, with WGS and RNA-Seq data as well as survival data available.
